# Biological Drivers of Early Childhood Caries in Preschool Children of Northern Arizona and Hawaii

**DOI:** 10.1101/2025.01.30.635340

**Authors:** Ryann N. Whealy, Tara N. Furstenau, Alexander Roberts, Jill Hager Cocking, Daryn Erickson, Breezy Brock, Rowan McCormick, Skylar Timm, Misty Pacheco, Summer Mochida-Meek, Viacheslav Y. Fofanov

**Affiliations:** Pathogen and Microbiome Institute, Northern Arizona University, Flagstaff, Arizona, United States of America; School of Informatics, Computing, and Cyber Systems, Northern Arizona University, Flagstaff, Arizona, United States of America; Department of Kinesiology and Exercise Sciences, University of Hawaiʻi at Hilo, Hilo, HI, United States of America

## Abstract

Early childhood caries (ECC) is the most prevalent chronic childhood disease, disproportionately affecting children from specific racial/ethnic groups. While ECC is a multifactorial disease influenced by demographic, behavioral, and environmental factors, it is also widely associated with the presence of *Streptococcus mutans* in the oral microbiome. To better understand the interplay between demographic and microbial factors and ECC risk, we collected saliva samples from 408 preschool children aged 1-6 years, including 266 from northern Arizona and 142 from Hawaii, representing racially and geographically diverse populations. Logistic regression showed that the odds of developing ECC increased 80.09% with each year of age, and that compared to White children, the odds were significantly higher for Native Hawaiian/Pacific Islander (330.94%), Native American (282.35%), and Hispanic (245.22%) children. Oral *S. mutans* colonization increased odds by 360.49%. The *S. mutans-*positive samples were genotyped using a multiplexed targeted amplicon sequencing assay. Phylogenetic analysis showed high genetic diversity in *S. mutans* between and within populations, with no geographic clustering. Some *S. mutans* clades were associated with up to a 33-fold increase in ECC odds. Only Native Hawaiian/Pacific Islander children were more likely to carry *S. mutans* strains from higher-risk clades. Markers associated with high risk *S. mutans* strains were identified in genes involved in carbohydrate metabolism, pH regulation, and biofilm formation. This study underscores the multifaceted nature of ECC, linking demographic disparities with strain-specific features of *S. mutans* and identifies new genetic markers that may play a role in *S. mutans* virulence.

## INTRODUCTION

Early childhood caries (ECC) is the most prevalent chronic childhood disease, affecting five times more children than asthma [1]. In the U.S., dental caries is found in the primary teeth of approximately 23% of children aged 2-5 years [2]. While ECC causes short term effects such as pain and difficulty eating, impacts may persist beyond preschool age, leading to missed school and general health effects [3,4]. Due to the chronic nature and delayed pain associated with dental caries, children are often not seen by a dental professional until it is too late to avoid severe symptoms and intensive treatment. This delay, in addition to gaps in dental insurance, means that treatment can often become a dental emergency; in 2012, $1.6 billion was spent on emergency dental visits [5]. Unfortunately, emergency room treatment for dental caries is typically aimed at reducing the pain and immediate infection, rather than resolving the underlying oral pathology, thus leading to return visits. Left untreated, the infection can spread through the fascial tissues to other parts of the head and neck, leading to hospitalization and, in rare cases, death [6].

Various measures of oral health show clear racial disparities in the United States. Children from socially disadvantaged backgrounds continue to have high rates of ECC, despite a substantial decrease in overall prevalence due to the significant progression of prevention and treatment strategies over recent decades [7,8]. Nationwide, caries rates are nearly doubled in Hispanic, Native American, and Native Hawaiian and Other Pacific Islander children compared to white children [2]. While this may be explained by restricted financial or physical access to care disproportionately affecting racial and ethnic minorities, additional barriers in access have been reported. Preferred language, education level, family income level, and nativity are all important determinants of oral health; intersecting identities often have additive effects, increasing the number of barriers faced [9]. Potentially related to these known oral health disparities, both Arizona and Hawai’i are historically among the worst states for dental health in the country, with rates of children impacted by tooth decay approximately double the national average [10–12]. Compared to the United States, Arizona’s population has higher proportions of American Indian and Hispanic/Latino individuals, and Hawaii has higher proportions of Asian, Native Hawaiian/ Pacific Islander, and multi-racial individuals. Additionally, Arizona has a higher-than-average percentage of the state living below the poverty line and without health insurance (12.4% and 12.1%, respectively), decreasing access to dental care [13].

In addition to the well-established socioeconomic risk factors for ECC, biological factors are thought to play an important role in the prevalence and severity of ECC. When compared to healthy oral plaque, pH in caries-associated plaque has been shown to remain low for abnormally extended periods of time after eating or drinking, thereby leading to demineralization of the tooth enamel [14]. In the oral cavity, excess carbohydrates, such as those in sugary drinks, potentially skew the oral microbiome towards creating more favorable conditions and thus select for increasingly pathogenic phenotypes. In fact, by lowering the pH of the oral cavity, cariogenic bacteria (which survives and thrives at lower pH levels) is thought to outcompete the normal “healthy” oral microbiota. Dental caries formation is most commonly associated with *Streptococcus mutans* [15–17]. This species is adapted to adhere to enamel and survive in the highly acidic environment created by the metabolism of readily fermented carbohydrates in the diet [18]. Thus, the amount of mutans streptococci in the oral cavity, virulence, strain identity, and the overall oral microbiota can all have a significant impact on the speed of caries progression and the efficacy of the intervention process.

The role that *S. mutans* genomic variation plays in ECC risk and how it impacts racial and ethnic disparities remains poorly understood. While traditional socioeconomic risk factors contribute to disparities in ECC incidence, they do not fully explain the elevated caries rates observed in certain racial and ethnic groups [19]. This gap suggests that biological factors, including strain-level genomic differences in *S. mutans,* may play a critical role in disease risk and progression. To investigate this, we focused on four key questions: 1) What are the differences in caries risk among the racial and ethnic demographics represented in Arizona and Hawaii? 2) Are *S. mutans* strains geographically distinct, or are they widely distributed across both locations? 3) Are specific *S. mutans* strains associated with higher caries risk, and are these strains linked to groups that are disproportionately impacted by caries? and 4) Can we identify potential genetic markers in *S. mutans* that are associated with caries risk?

## METHODS

### Study participants

Children aged 1-5 years from 14 participating preschool and daycare locations in Coconino County, Arizona were recruited for sample collection in addition to free dental care services provided by Smart Smiles, a school-based oral health program. Four sampling campaigns, lasting six months each, were completed between Fall 2017 and Spring 2019. Additionally, children aged 1-6 and their parents were recruited from five participating preschool locations in Hawaii for sample collection by the University of Hawaiʻi at Hilo. Two sampling campaigns were completed between Fall 2021 and Spring 2022. School administrators opted into the study, and parents independently consented to dental care services and sample collection. Parents were made aware that bacterial genomics were the focus of this study and no human genetic material would be sequenced. Parents provided demographic information for their child, including age, gender, and ethnicity. The number and extent of caries affected teeth were provided by Smart Smiles or University of Hawaiʻi at Hilo during health screenings.

### Sample Collection

BD ESwab collection and transport system was used to collect saliva samples, using the standard, minimally invasive, and widely-used microbiome protocol of the Human Microbiome Project (HMP). Specifically, non-stimulated saliva was collected by swabbing the buccal mucosa with a sterile cotton swab. Collection was carried out by a trained dental hygienist to minimize risk to the child. Samples were stored on dry ice in 1 mL of modified liquid Amies medium during transport and then at -80°C until processing.

### qPCR detection of *S. mutans*

DNA was extracted from collected colonies using the Zymo Quick-DNA Fungal/Bacterial Miniprep Kit. The presence of *S. mutans* was confirmed using a previously established qPCR assay [20]. The qPCR included TaqMan Universal Master Mix with primers at a final concentration of 0.6 µM and probes at 0.25 µM. The optimized cycle conditions for this reaction was: 50°C for 2 minutes, 95°C for 10 minutes, and 50 cycles of 95°C for 15 seconds and 60°C for 1 minute.

### Development of AmpSeq Assay

A targeted amplicon sequencing assay was developed for high-resolution genotyping the *S. aureus* samples without requiring whole genome sequencing for all samples. The amplicon targets were chosen using VaST v1.0.0 [21] which identifies a minimal set of target loci that maximize differentiation between genomes. The loci were chosen among 81,473 core genome SNPs discovered from an alignment of 190 *S. mutans* genomes downloaded from NCBI complete genomes database (all assembly levels up to the year 2018) [22] using NASP v1.2.1 [23] with NC_004350.2 as a reference (Supplemental Table 1). A total of 29 amplicon targets (covering 500 of the SNPs discovered from the reference sequences) were selected for the panel and provided high genotype resolution of the reference strains. Supplemental Figure 1 shows the minimum spanning tree drawn with GrapeTree [24]. The average amplicon size (without primer regions) was 220bp (min=89, max=387) and the total size of all amplicons was 6,834. Primers were designed to amplify these targets in regions that were highly conserved across the reference genomes and the primers were optimized to work together in a single multiplex PCR (Supplemental Table 2).

### Molecular Characterization

#### Whole genome sequencing

Whole genome sequencing (WGS) was performed on a subset of *S. mutans-*positive samples (qPCR) from the northern Arizona cohort to compare our AmpSeq assay to WGS. These samples were streaked onto TYCSB media using the original swab and a plastic inoculating loop followed by incubation for 48 hours at 37°C. A single *S. mutans* colony, identified based on morphology, was isolated from each plate, and DNA was extracted using the Zymo Quick-DNA Fungal/Bacterial Miniprep Kit. Samples were sequenced on an Illumina MiSeq instrument to produce 300bp paired end reads. To compare the phylogenetic placement of samples that underwent parallel WGS and AmpSeq, we used WG-FAST [25] to infer a maximum likelihood tree using SNPs called from these samples and SNPs identified in the 190 reference genomes. Supplemental Figure 2 shows that both the WGS and AmpSeq samples displayed similar placement in the tree, suggesting that the phylogenetic signal captured by our AmpSeq targets is strong and consistent with the WGS-derived data.

#### Amplicon sequencing

DNA was extracted directly from the transport media for all *S. mutans-*positive samples (qPCR) using the Zymo Quick-DNA Fungal/Bacterial Miniprep Kit. For the multiplex PCR, The KAPA 2G Fast Multiplex PCR Master Mix was used with 29 *S. mutans-*specific primer pairs (Supplemental Table 2). Primers were diluted to a final concentration of 0.2 *µ*M. The optimized cycle profile was: 95°C for 3 minutes, 35 cycles of 95°C for 15 seconds, 60°C for 30 seconds, and 72°C for 1 minute 30 seconds, 72°C for 1 minute, and 10°C indefinitely. Successful amplification was confirmed via gel electrophoresis, followed by a bead cleanup using a 2:1 ratio of AMPure (Beckman Coulter) beads. Barcodes were added through an extension PCR with the following cycle profile: 98°C for 2 minutes, 6 cycles of (98°C for 30 seconds, 60°C for 20 seconds, and 72°C for 30 seconds), 72°C for 5 minutes, and 10°C indefinitely. A second bead clean up was performed with the same conditions. Libraries were quantified for pooling using a KAPA quantification kit, and the pooled libraries underwent fragment analysis for final quality assurance. Samples were sequenced on an Illumina MiSeq Platform.

### Sequence processing and analysis

Reads from both WGS and AmpSeq were quality trimmed and filtered using Fastp v0.20.1 [26] with default parameters. The sequences were aligned to reference *S. mutans* UA159 (NC_004350.2) using BWA-MEM v0.7.8 [27] and SNPs were called using UnifiedGenotyper v3.4.46 [28]. SNPs within the expected amplicon regions (excluding primer regions) were concatenated and used to build a maximum likelihood tree with IQTree v1.6.12 [29] after filtering samples with less than 50% callable sites. IQTree was run with the best-fit model GTR+F+G4. There were a total of 173 sequences in the alignment and 250 SNPs (216 parsimony-informative and 34 singletons).

### Classifying *S. mutans* genotypes into caries risk groups using K-nearest neighbors

*S. mutans* genotypes were classified into caries risk groups using a K-nearest neighbors approach based on tree distance. A distance matrix was created from the maximum likelihood tree using Dendropy v5.0.1 [30]. For each *S. mutans* sample, the proportion of caries-positive samples from the 5 nearest samples (or more if there was a tie for the same distance) was calculated (multiple samples from the same individual were only counted once). We considered a caries-positive proportion less than 25% to be low risk, from 25%-50% to be medium risk and greater than 50% to be high risk.

### Statistical analysis

Cavities were treated as a binary variable, where individuals with at least one cavity were coded as 1, and those with no cavities were coded as 0. Race and ethnicity were categorized using indicator variables for specific groups, including Native American, Black, Native Hawaiian/Pacific Islander, Hispanic, Asian, and Non-Hispanic White. For all models, Non-Hispanic White was used as the reference group. Risk categories (low, medium, high) were numerically re-coded as 0, 1, and 2, respectively. For clade analysis, the reference group was set to Clade 161, which was exclusively observed in children in the low risk category. *S. mutans* colonization was coded as 1 if the individual had positive results using qPCR or AMPSeq, and 0 otherwise. Participant metadata and classifications are provided in Supplemental Table 3.

Logistic regression was used to identify factors associated with presence of at least one cavity. The full model included age, race/ethnicity, sex, and *S. mutans* colonization; predictors were individually removed according to significance. Additionally, a penalized logistic regression was used to evaluate the association between clade membership and cavity presence, addressing separation in the data due to varied clade sizes. Logistic regression models were evaluated using receiver operating characteristic curves and AIC. Finally, a generalized linear model was used to assess the impact of demographic factors on risk categories. The full model included age, race/ethnicity, and sex.

### Correlation between geographical regions and phylogenetic placement

We used a mantel test to estimate the strength of the associate between the two different collection regions (Hawaii vs. Arizona) and the placement of the samples in the phylogenetic tree. The Mantel test was performed using Scikit-Bio v0.6.2 [31] using Spearman correlation with 1,000 permutations on the genetic distance matrix from the maximum likelihood tree and a binary spatial distance matrix.

### *S. mutans* genetic markers associated with caries risk

To identify genome wide markers and genes associated with caries risk, we used our k-nearest neighbors classification approach on a collection of 190 *S. mutans* reference genomes and identified significant associations with predicted caries risk and the presence or absence of genes and SNPs. To predict the caries risk for the reference genomes, a maximum likelihood tree was inferred using IQ-Tree from concatenated SNPs in the targeted amplicon regions from our samples and the reference genomes. There were a total of 467 SNPs (340 parsimony-informative, 127 singletons) and the best-fit model was SYM+ASC+G4. In the same way as described above, the 5 nearest samples (based on genetic distance) were selected for each reference sequence and the proportion of samples with caries was used as the risk score. A table of core genome SNPs was created for the reference genomes as described above and contained 81,472 SNPs. To identify the presence or absence of genes, the genomes were consistently annotated using Prokka v1.14.6 [32] and Roary v3.13.0 [33] was used to define the core and pan genome. Out of a total of 9,806 genes considered, there were 788 core genes (in 99% or more of strains), 469 soft core genes (in 95-99% of strains), 867 shell genes (in 15-95% of strains), and 7,683 cloud genes (in 0-15% of strains).

We analyzed the results using two different approaches that differed in whether they took population structure into account. First we used XGBoost, an implementation of gradient boosted decision trees, to identify SNPs predictive of caries risk scores. The core genome SNP table (encoded as binary variables) and the predicted caries risk scores were analyzed using the XGBRegressor module from the XGBoost Python package v2.1.3 [34] trained using 100 estimators. To evaluate the contribution of individual SNPs to the prediction of risk score, Shapley additive explanation (SHAP) values were computed using the ‘shap’ Python library v0.46.0 [35]. SNPs were ranked based on their mean absolute SHAP values, and the top 20 SNPs were annotated. This approach did not account for population structure inherent in the tree and therefore will identify SNPs that are strongly associated with high and low risk clades but the association may be due to shared ancestry rather than a true causal relationship.

For our second approach, we analyzed the core genome SNPs and the presence/absence of 9,806 genes using Pyseer v1.3.12 [36]. Pyseer does account for shared ancestry between the genomes so the features identified using this tool are more likely to be causally linked to caries risk. The population structure was provided by providing a distance matrix that was created from the phylogenetic tree using Dendropy v5.0.1 [37]. Genes and SNPs were considered to be significantly associated with predicted caries risk if their p-value was below the threshold (p<0.05) with a Bonferroni correction applied for multiple testing.

## RESULTS

### Sample Collection and Sequencing

We collected samples from 408 children, 266 from northern Arizona and 142 from Hawaii. We detected and sequenced *S. mutans* from 202 samples from 167 children (99 from northern Arizona and 68 from Hawaii) for a total of 217 sequences. The full sample metadata is provided in Supplemental Table 3.

### Risk factors for caries development

Logistic regression was used to evaluate the risk factors associated with the presence of at least one cavity while adjusting for all other variables. The analysis showed that the odds of having at least one cavity increased by 80.09% for each additional year of age (*p* < .001). Significant differences in cavity risk were also observed across racial and ethnic groups. Compared to White children, the odds were 714.20% higher for Black children (*p* = 0.020), 330.94% higher for Native Hawaiian/Pacific Islander children (*p* < .001), 282.35% higher for Native American children (*p* = 0.006), 345.22% higher for Hispanic children (*p* = 0.005), and 214.68% higher for Asian children (*p* = 0.005). Additionally, the odds of having at least one cavity were 360.49% higher for children with oral *S. mutans* colonization (*p* < .001).

### *S. mutants* genotypes are not geographically constrained

The phylogenetic relationships of *S. mutans* collected from the two geographical regions (Northern Arizona and Hawaii) did not show any strong clustering by geographical region. Genotypes from both locations were interspersed throughout the phylogenetic tree rather than clustering into region-specific clades (Figure 1). The Mantel correlation coefficient was r = 0.046 (*p* = 0.011) indicating weak association between phylogenetic and geographical distance. These results suggest that the *S. mutans* strains exhibit substantial genetic diversity within and between populations and they are widely distributed.

**Figure 1.**
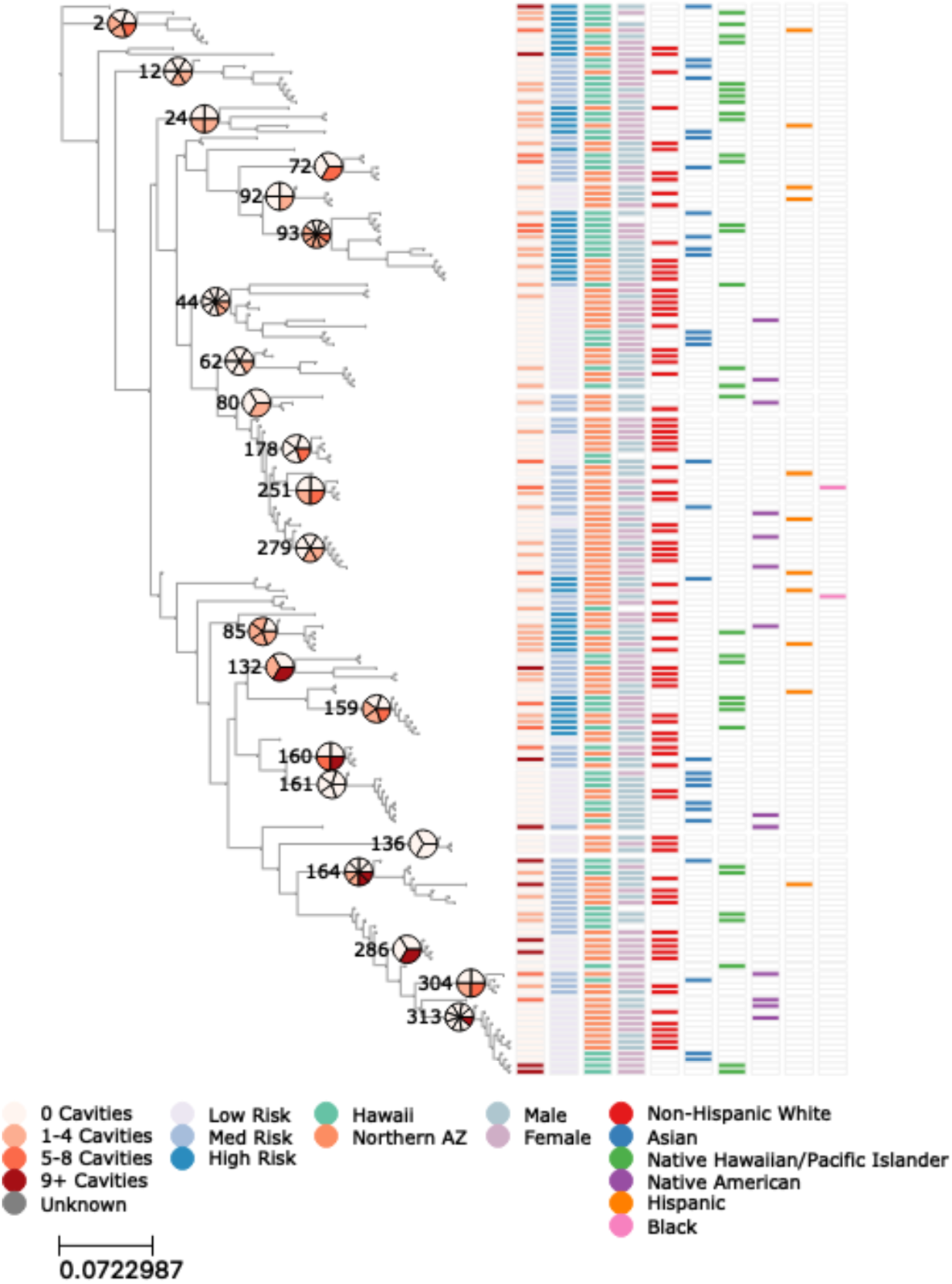
*S. mutants* clades exhibit differences in caries risk. The maximum likelihood tree of *S. mutans* strains from positive samples shows the number of cavities reported for the child (column 1) and the estimated risk based on the caries status of K-nearest neighbors in the tree (column 2). There is clear clustering of high, medium, and low caries risk *S. mutans* strains within clades. No noticeable clustering was observed for location (column 3), sex (column 4), or reported race and ethnicity (columns 5-10). The pie charts indicate the caries status of children associated with the *S. mutans* strains at select nodes in the tree.

### *S. mutans* clade membership is a significant predictor of caries

Firth’s penalized logistic regression was used to quantify the significance of *S. mutans* clades in their predictive value of cavity presence. This model had an AUC of 0.76, indicating good discriminative ability. Compared to the reference clade, which was only identified in children with low risk of cavities, odds of having at least one cavity were increased by a factor of: 15.4 in clade 2 (*p* = 0.055), 18.3 in clade 24 (*p* = 0.062), 33.0 in clade 85 (*p* = 0.013), 15.9 in clade 93 (*p* = 0.030), 18.3 in clade 132 (*p* = 0.062), 18.3 in clade 160 (*p* = 0.062), and 11.0 in clade 164 (*p* = 0.075).

A generalized linear model was used to examine the impact of demographic features on the likelihood of belonging to a low or high-risk clade. Native Hawaiian/Pacific Islander children were the only group significantly different from white children; on a risk category scale of 0-2 (low, medium, high), NH/PI children had an average risk category that was 0.46 points higher than that of white children (*p* = 0.004), with an expected risk category value of 1.27. In contrast, sex and age were not found to be significant predictors of clade membership.

### *S. mutans* markers associated with caries risk

To identify genetic markers predictive of caries risk, we employed two complementary approaches: XGBoost and Pyseer. XGBoost, which does not account for population structure, was used to identify SNPs that strongly correlate with the caries risk predictions made for the reference genomes. This approach had the advantage of highlighting associations that might drive clustering in the phylogenetic tree. However, it may also detect SNPs that are associated with risk due to shared ancestry rather than true causal relationships. In contrast, Pyseer accounted for population structure by incorporating a phylogenetic distance matrix, providing a framework to identify SNPs and genes more likely to have a direct causal role in caries risk.

With XGBoost, the top 20 core genome SNPs with the highest feature importance, as determined by SHAP values (Supplemental Figures 3 and 4), were identified. A majority of these SNPs (15/20) were located in coding regions (Table 2), while most of the non-coding SNPs (4/5) were situated directly upstream or downstream of a coding region (4/5). Only one of the SNPs produced a non-synonymous mutation and this was in a multiple sugar transport system substrate-binding protein. When these coding regions were compared to our previous metatranscriptomic study of paired cavity and non-cavity plaque samples, 13 were found to be significantly upregulated in cavity plaque, suggesting a potential functional role in caries [38].

**Table 1.** Demographics of participants.

| Group | Category | HI | NAZ | Total |
| --- | --- | --- | --- | --- |
| Gender | Girl | 66 | 128 | 194 |
|  | Boy | 52 | 136 | 188 |
|  | Unknown | 24 | 2 | 26 |
| Age | 1 | 0 | 30 | 30 |
|  | 2 | 7 | 45 | 52 |
|  | 3 | 37 | 73 | 110 |
|  | 4 | 55 | 81 | 136 |
|  | 5 | 19 | 33 | 52 |
|  | 6 | 1 | 0 | 1 |
|  | Unknown | 23 | 4 | 27 |
| Race/ Ethnicity | Non-Hispanic White | 8 | 191 | 199 |
|  | Asian | 50 | 4 | 54 |
|  | Black | 2 | 6 | 8 |
|  | Hispanic | 1 | 25 | 26 |
|  | Native American | 0 | 35 | 35 |
|  | Native Hawaiian/ Pacific Islander | 57 | 1 | 58 |
|  | Unknown | 24 | 4 | 28 |

**Table 2.**
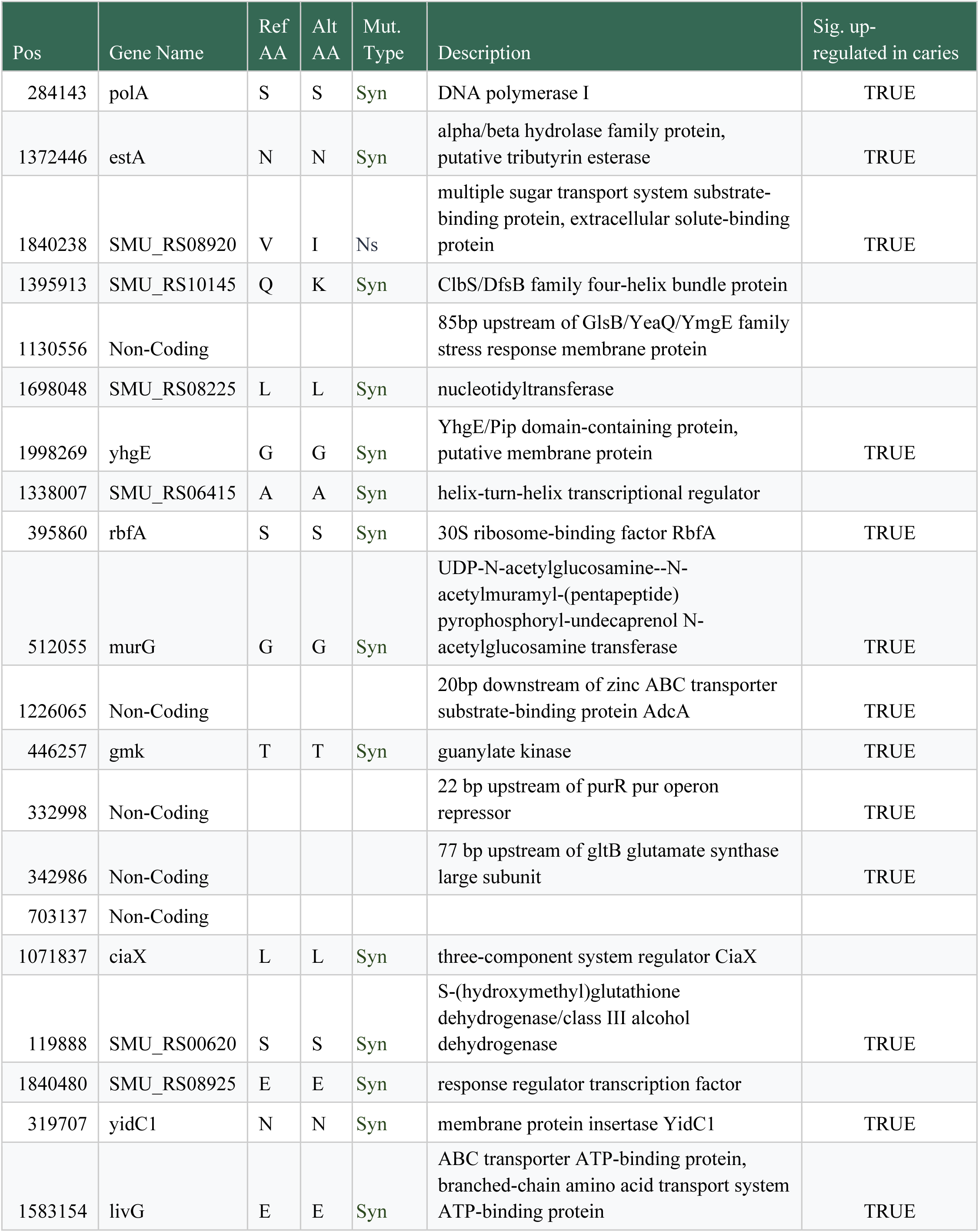
SNP annotations with for top 20 highest feature importance identified by XGBoost. The SNPs are ordered from most to least important. SNPs in genes that were previously determined to be significantly upregulated in caries in our metatranscriptomic study are identified in the last column.

In our analysis of genes using PySeer, we identified significant associations for multiple genetic features. We observed a strong negative association (ꞵ = -0.273) for a gene annotated as *lagD*, which codes for a Lactococcin-G-processing and transport ATP-binding protein, and was present in 8 of the genomes. This protein plays a critical role in the production and export of lactococcin G, a bacteriocin with antimicrobial properties [39]. The strong negative association suggests that the presence of this gene may reduce caries risk by modulating the microbial community. We also identified a strong negative association for a gene annotated as a helix-turn-helix-type transcriptional regulator CysL (ꞵ = -0.2, present in 166 of the genomes) which is involved in sulfur metabolism with links to biofilm formation, oxidative stress resistance, and microbial interactions [40].

There were 165 core genome SNPs that PySeer identified as significant (Supplemental Table 4), 46% (mean ꞵ = 0.31) were positively associated with caries risk and 54% (mean ꞵ = -0.27) were negatively associated. Most of the significant SNPs were in coding regions (87%) and 36% were non-synonymous. 60% of the SNPs were in genes that we previously found to be significantly upregulated in caries, and 41% of these were non-synonymous. SNPs in the same AraC-family helix-turn-helix transcriptional regulator gene (SMU_RS06415) were identified using both PySeer and XGBoost but they were at different positions. Other notable genes with significant non-synonymous SNPs include: *argG* (argininosuccinate synthase), an enzyme that is part of arginine metabolism and plays a role in maintaining oral pH, *nagA*, an enzyme that plays a role in carbohydrate metabolism, *gltB*, an enzyme involved in glutamate metabolism that contributes to stress tolerance and pH regulation, *ftsY* a fused signal recognition particle receptor related to biofilm formation, *recN*, a DNA repair protein important for survival through oxidative stress and in challenging environments, and *pepX*, *pepF*, and *pepB*, enzymes involved in peptide metabolism.

## DISCUSSION

This study investigated the demographic and microbial genomic factors associated with ECC risk. Age, race/ethnicity, and oral *S. mutans* colonization emerged as significant predictors of ECC in this study. The strong association between increasing age and caries risk (80.09%, *p* < 0.0005) aligns with previous research, emphasizing the cumulative effects of prolonged exposure to cariogenic factors and the critical need for early preventative interventions [41]. We found that Native American, Native Hawaiian/Pacific Islander, and Hispanic children were at a significantly higher risk for caries development than White children (Black children were also found to have a significantly higher risk of caries development; however, the small sample size limits the statistical power to confidently support this finding). These disparities in risk likely reflect the socioeconomic factors contributing to caries development, including dietary patterns, access to preventative care, and oral health literacy, as various existing studies have demonstrated [42].

Our results confirm that *S. mutans* is a significant risk factor for ECC. Phylogenetic analysis of 173 sequenced *S. mutans* strains revealed wide genetic diversity, with over 250 SNPs identified across the 6,834 total base pairs targeted by PCR. We observed no significant geographic clustering of genotypes between Arizona and Hawaii, aligning with previous findings that *S. mutans* strains are widely distributed geographically [43]. This lack of geographic specificity has important epidemiological implications as it suggests that high-risk *S. mutans* strains are not confined to localized populations but are broadly distributed.

Certain *S. mutans* clades were associated with varying levels of caries risk. For example, clade 85 was associated with a 33-fold increase in the odds of having cavities (*p* = 0.013). Clades 2, 24, 93, 132, 160, and 162 displayed strong predictive values as well, though at varying levels of significance. Native Hawaiian and Pacific Islander children had significantly increased odds of carrying *S. mutans* strains from high-risk clades. This suggests that some of the disparities in ECC observed in this population may be partially influenced by unique host-strain dynamics. Future research should investigate if this is due to socioeconomic, behavioral, environmental, or other biological factors.

We performed a pseudo-genome-wide association study to identify genetic predictors of caries risk. Because WGS data was not available for all samples, we used the placement of publicly available complete reference genomes within the phylogenetic tree of our samples as a proxy. Caries risk for these genomes was estimated using k-nearest neighbors classification based on the genetic distance with our samples. This approach enabled us to identify genome-wide features associated with caries risk, including both the presence/absence of genes and core genome SNPs. Many of the markers identified were linked to genes involved in critical processes such as carbohydrate metabolism, signalling, pH regulation, biofilm formation, and survival within the oral environment. While our approach was only an approximation of a traditional genome wide association study, its results are supported by our previous metatranscriptomic study [38]. Notably, 90% of the coding regions associated with significant markers were identified in the previous study as being significantly upregulated in caries-active plaque compared to non-caries plaque. This overlap suggests a functional role for these markers in caries pathogenesis, warranting further validation in future studies.

## CONCLUSIONS

This study highlights the multifactorial nature of ECC, demonstrating that age, race/ethnicity, and microbial genetic factors collectively influence caries risk. Disparities among racial and ethnic groups likely arise from a complex interplay between social determinants and biological susceptibility. The identification of high-risk *S. mutans* clades and genetic markers provides a foundation for strain-specific diagnostics and targeted interventions. These findings underscore the need for integrated public health strategies that account for microbial diversity in addition to other risk factors to effectively reduce ECC disparities in young children.

---

**Supplemental Figure 1.**
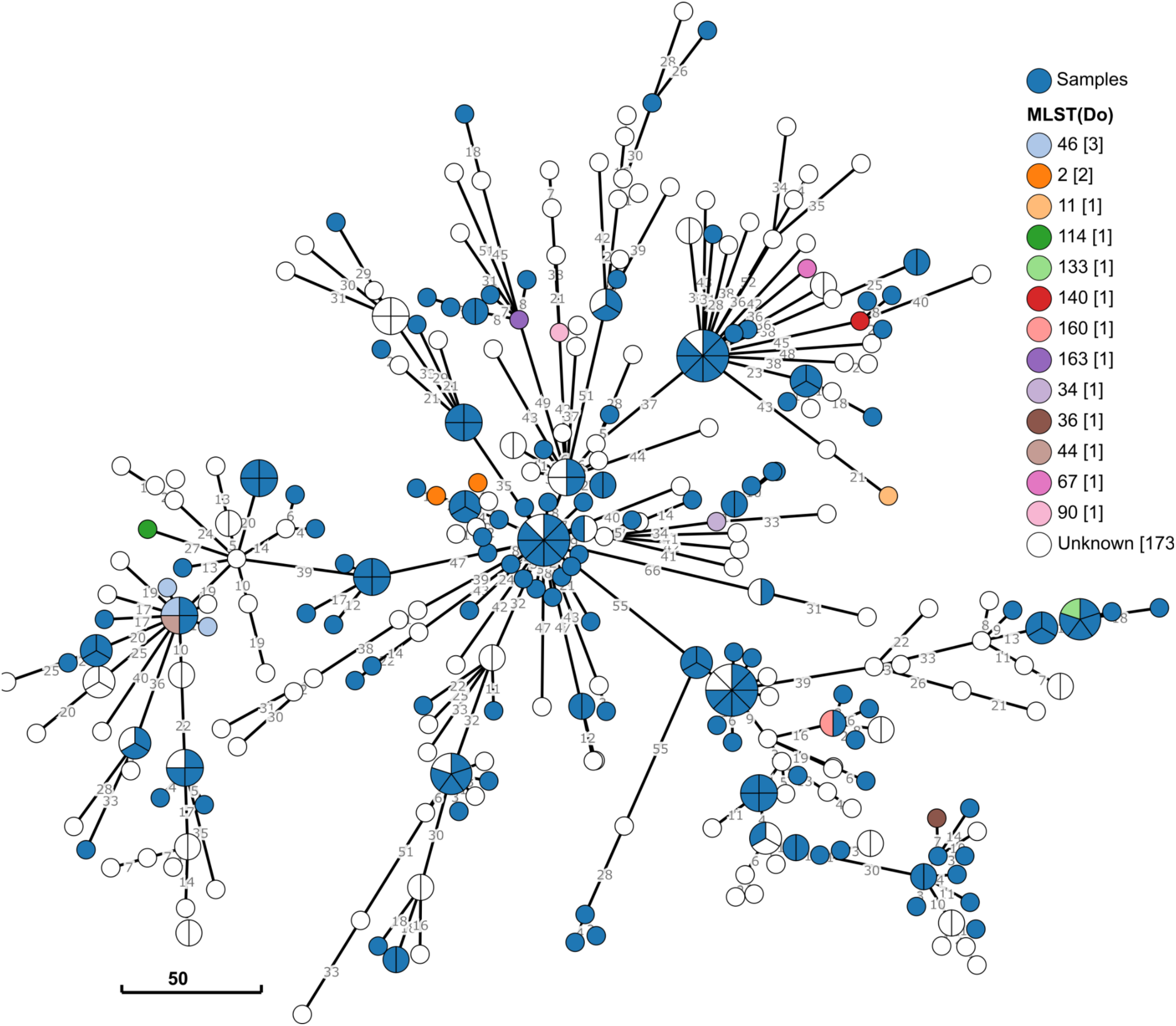
Minimum spanning tree of SNPs from selected amplicon targets showing the differentiation of 190 reference strains used to design the amplicon sequencing panel and samples sequenced for this project. The branch lengths are indicated and the nodes are pie charts showing the number of unresolved genomes at each node. Where MLST typing (Do et al, 2009) was possible for the reference genomes, the sequence type are color coded.

**Supplemental Figure 2.**
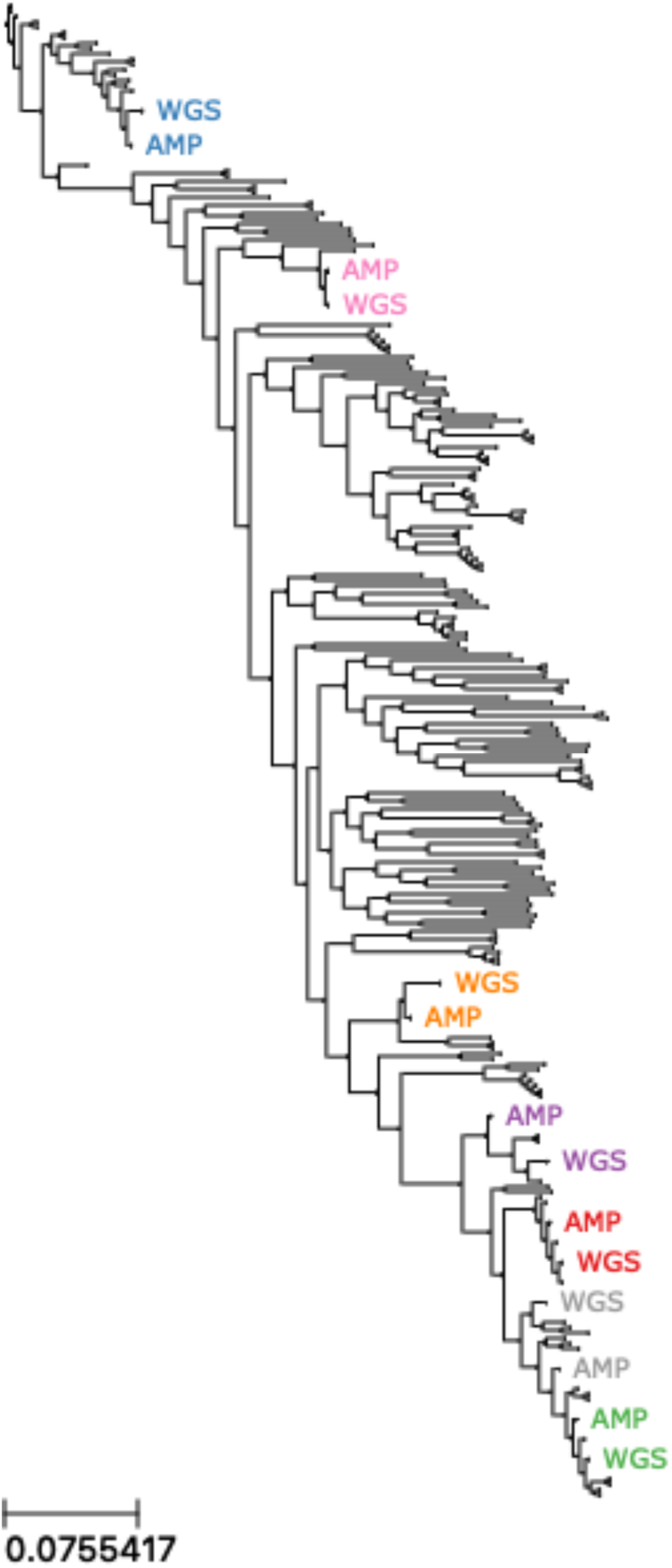
Samples analyzed using both AmpSeq and WGS sequencing show similar placement in maximum likelihood phylogeny. The maximum likelihood phylogeny was inferred using SNPs called from samples that underwent parallel AmpSeq and WGS along with SNPs from 190 reference genomes. The colors indicate the different samples and the sequencing type is indicated. In each case, both samples are within the same clade.

**Supplemental Figure 3.**
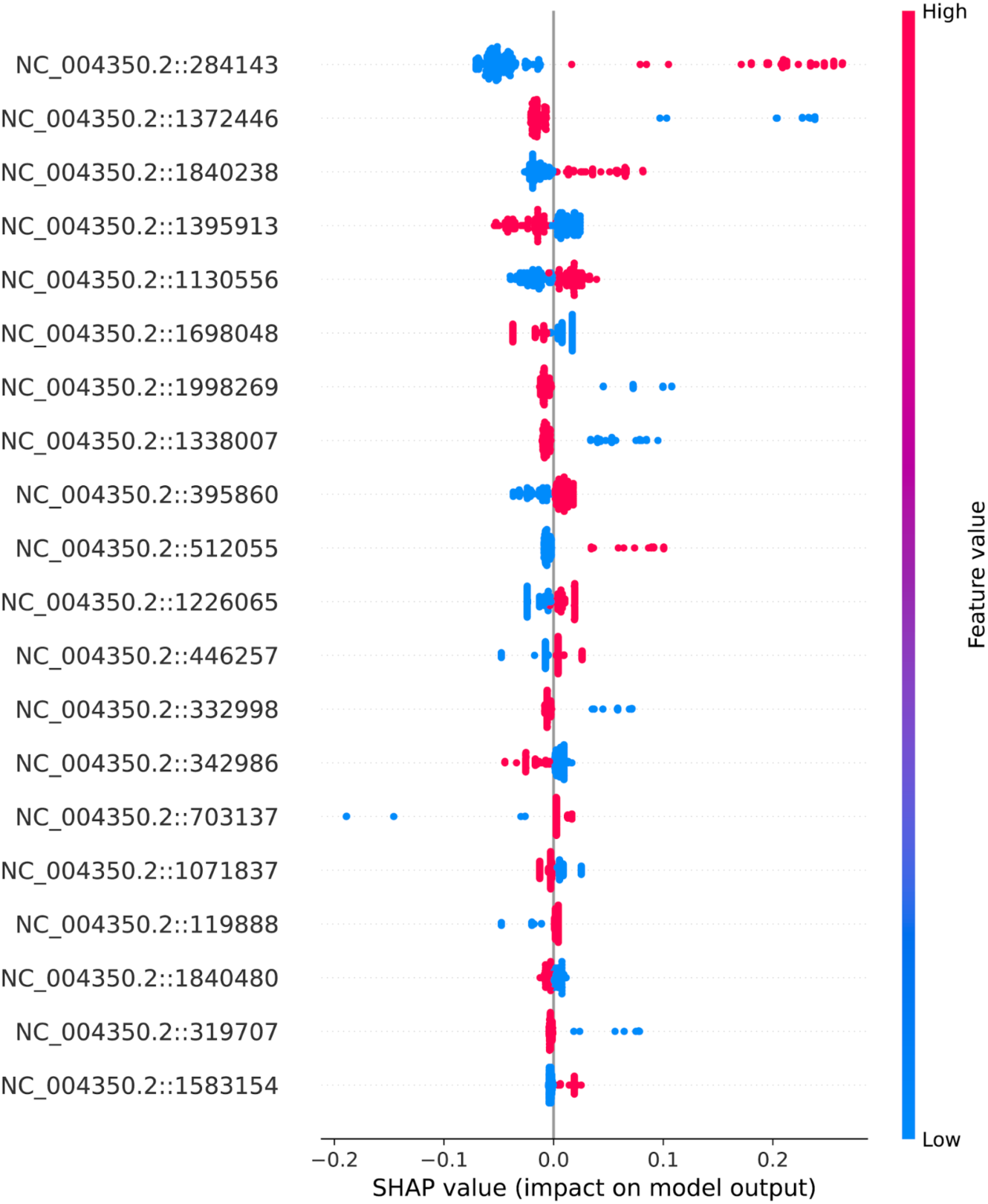
SHAP values for the top 20 SNPs identified through XGBoost. SNPs are ordered by importance (highest to lowest). Each point represents a SHAP value for an individual SNP, with blue and red points denoting the binary states of the SNP. Positive SHPA values indicate that the SNP increased the predictive outcome (i.e., higher risk), while negative SHAP values suggest a decrease in the predictive outcome. SNPs with SHAP values further from zero and with larger separation between the binary states demonstrate greater feature importance.

**Supplemental Figure 4.**
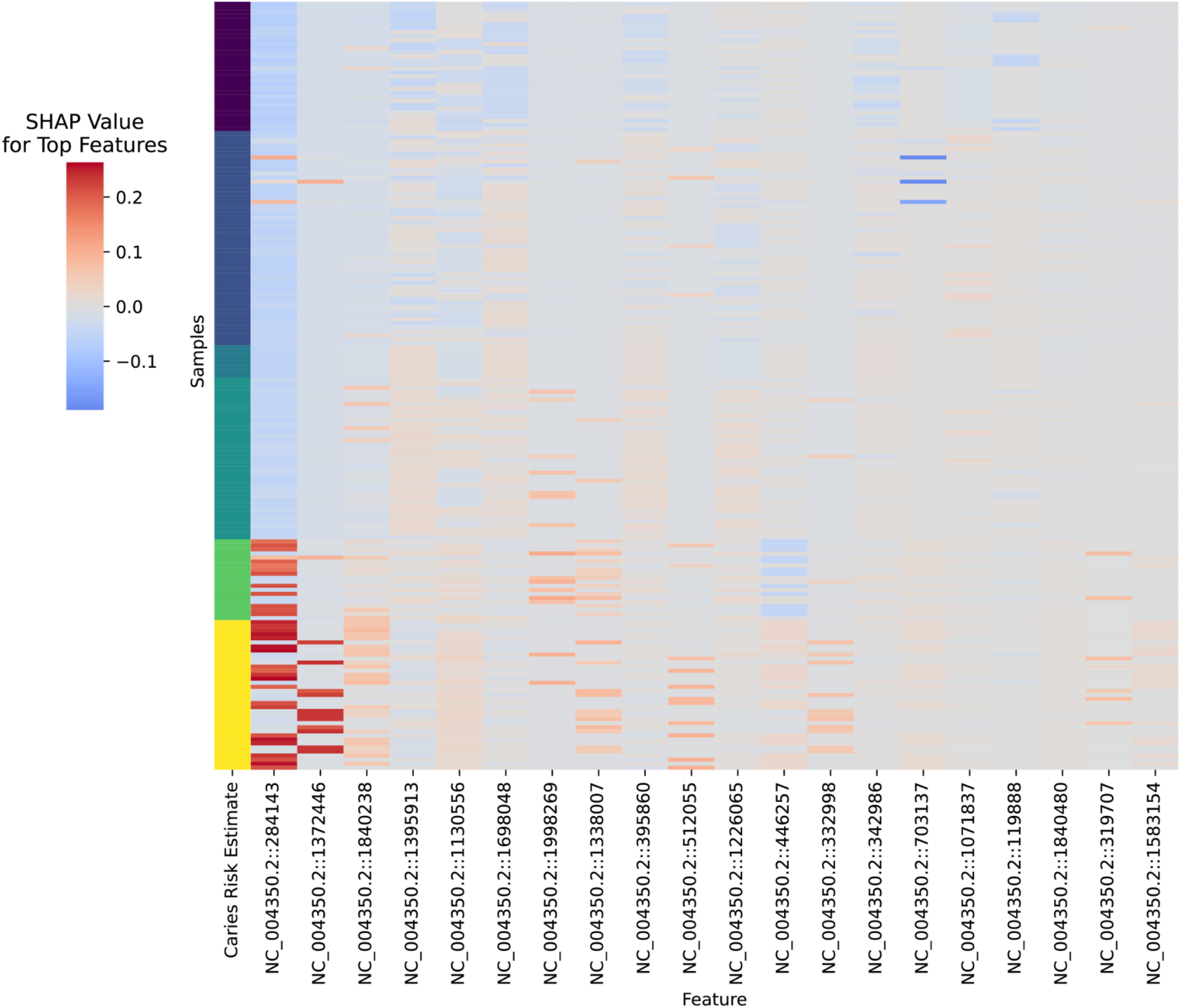
Heatmap of SHAP values for top 20 SNPs and associated caries risk estimate for each sample. Rows represent individual samples and columns correspond to the top 20 SNPs identified with XGBoost (in order of importance from left to right). The color indicates the magnitude and direction of the SHAP values, with positive values contributing to higher caries risk estimates and negative values contributing to lower risk. The column on the left indicates the caries risk estimates provided as the outcome variables to the model.

**Supplemental Table 1:** Reference sequence assembly accessions.

| Reference Assembly Accessions |
| --- |
| GCF_000007465.2 |
| GCF_000091645.1 |
| GCF_000228745.1 |
| GCF_000228765.1 |
| GCF_000228785.1 |
| GCF_000228805.1 |
| GCF_000228825.1 |
| GCF_000228845.1 |
| GCF_000228865.1 |
| GCF_000228885.1 |
| GCF_000228905.1 |
| GCF_000228925.1 |
| GCF_000228945.1 |
| GCF_000228965.1 |
| GCF_000228985.1 |
| GCF_000229005.1 |
| GCF_000229025.1 |
| GCF_000229045.1 |
| GCF_000229065.1 |
| GCF_000229085.1 |
| GCF_000229105.1 |
| GCF_000229125.1 |
| GCF_000229145.1 |
| GCF_000229165.1 |
| GCF_000229185.1 |
| GCF_000229205.1 |
| GCF_000229225.1 |
| GCF_000229245.1 |
| GCF_000229265.1 |
| GCF_000229285.1 |
| GCF_000229305.1 |
| GCF_000229325.1 |
| GCF_000229345.1 |
| GCF_000229365.1 |
| GCF_000229385.1 |
| GCF_000229405.1 |
| GCF_000229425.1 |
| GCF_000229445.1 |
| GCF_000229465.1 |
| GCF_000229485.1 |
| GCF_000229505.1 |
| GCF_000229525.1 |
| GCF_000229545.1 |
| GCF_000229565.1 |
| GCF_000229585.1 |
| GCF_000229605.1 |
| GCF_000229625.1 |
| GCF_000229645.1 |
| GCF_000229665.1 |
| GCF_000229685.1 |
| GCF_000229705.1 |
| GCF_000229725.1 |
| GCF_000229745.1 |
| GCF_000229765.1 |
| GCF_000229785.1 |
| GCF_000229805.1 |
| GCF_000229825.1 |
| GCF_000229845.1 |
| GCF_000229865.1 |
| GCF_000229885.1 |
| GCF_000229905.1 |
| GCF_000229925.1 |
| GCF_000229945.1 |
| GCF_000229965.1 |
| GCF_000229985.1 |
| GCF_000230005.1 |
| GCF_000230025.1 |
| GCF_000230045.1 |
| GCF_000230065.1 |
| GCF_000230085.1 |
| GCF_000230105.1 |
| GCF_000230125.1 |
| GCF_000230145.1 |
| GCF_000230165.1 |
| GCF_000230185.1 |
| GCF_000230205.1 |
| GCF_000230225.1 |
| GCF_000271865.1 |
| GCF_000284575.1 |
| GCF_000339055.1 |
| GCF_000339075.1 |
| GCF_000339095.1 |
| GCF_000339115.1 |
| GCF_000339135.1 |
| GCF_000339155.1 |
| GCF_000339175.1 |
| GCF_000339195.1 |
| GCF_000339215.1 |
| GCF_000339235.1 |
| GCF_000339255.1 |
| GCF_000339275.1 |
| GCF_000339295.1 |
| GCF_000339315.1 |
| GCF_000339335.1 |
| GCF_000339355.1 |
| GCF_000339375.1 |
| GCF_000339395.1 |
| GCF_000339415.1 |
| GCF_000339435.1 |
| GCF_000339455.1 |
| GCF_000339475.1 |
| GCF_000339495.1 |
| GCF_000339515.1 |
| GCF_000339535.1 |
| GCF_000339555.1 |
| GCF_000339575.1 |
| GCF_000339595.1 |
| GCF_000339615.1 |
| GCF_000339635.1 |
| GCF_000339655.1 |
| GCF_000339675.1 |
| GCF_000339695.1 |
| GCF_000339715.1 |
| GCF_000339735.1 |
| GCF_000339755.1 |
| GCF_000339775.1 |
| GCF_000339795.1 |
| GCF_000339815.1 |
| GCF_000339835.1 |
| GCF_000339855.1 |
| GCF_000339875.1 |
| GCF_000339895.1 |
| GCF_000339915.1 |
| GCF_000339935.1 |
| GCF_000339955.1 |
| GCF_000339975.1 |
| GCF_000339995.1 |
| GCF_000340015.1 |
| GCF_000340035.1 |
| GCF_000340055.1 |
| GCF_000340075.1 |
| GCF_000340095.1 |
| GCF_000340115.1 |
| GCF_000340135.1 |
| GCF_000340155.1 |
| GCF_000340175.1 |
| GCF_000347795.1 |
| GCF_000347815.1 |
| GCF_000347835.1 |
| GCF_000347855.1 |
| GCF_000347875.1 |
| GCF_000347895.1 |
| GCF_000375505.1 |
| GCF_000496535.1 |
| GCF_000496555.1 |
| GCF_000522565.1 |
| GCF_000522585.1 |
| GCF_000522605.1 |
| GCF_000522625.1 |
| GCF_000522645.1 |
| GCF_000522665.1 |
| GCF_000522685.1 |
| GCF_000522705.1 |
| GCF_000522725.1 |
| GCF_000522745.1 |
| GCF_000522765.1 |
| GCF_000522785.1 |
| GCF_000522805.2 |
| GCF_000522825.1 |
| GCF_000522845.1 |
| GCF_000522865.1 |
| GCF_000522885.1 |
| GCF_000522905.1 |
| GCF_000522925.1 |
| GCF_000522945.1 |
| GCF_000817065.1 |
| GCF_001068415.1 |
| GCF_001069835.1 |
| GCF_001073145.1 |
| GCF_001558215.1 |
| GCF_001625005.1 |
| GCF_001703615.1 |
| GCF_002083175.2 |
| GCF_002155285.1 |
| GCF_002157665.1 |
| GCF_002179995.1 |
| GCF_002212845.1 |
| GCF_002212855.1 |
| GCF_002212885.1 |
| GCF_002212905.1 |
| GCF_002212925.1 |
| GCF_002212935.1 |
| GCF_002212965.1 |
| GCF_002212995.1 |
| GCF_002213005.1 |
| GCF_002213035.1 |
| GCF_002213065.1 |
| GCF_002995555.1 |
| GCF_003466855.1 |
| GCF_003691695.1 |
| GCF_900459345.1 |
| GCF_900475095.1 |
| GCF_900636835.1 |
| GCF_900638045.1 |

**Supplemental Table 2:**
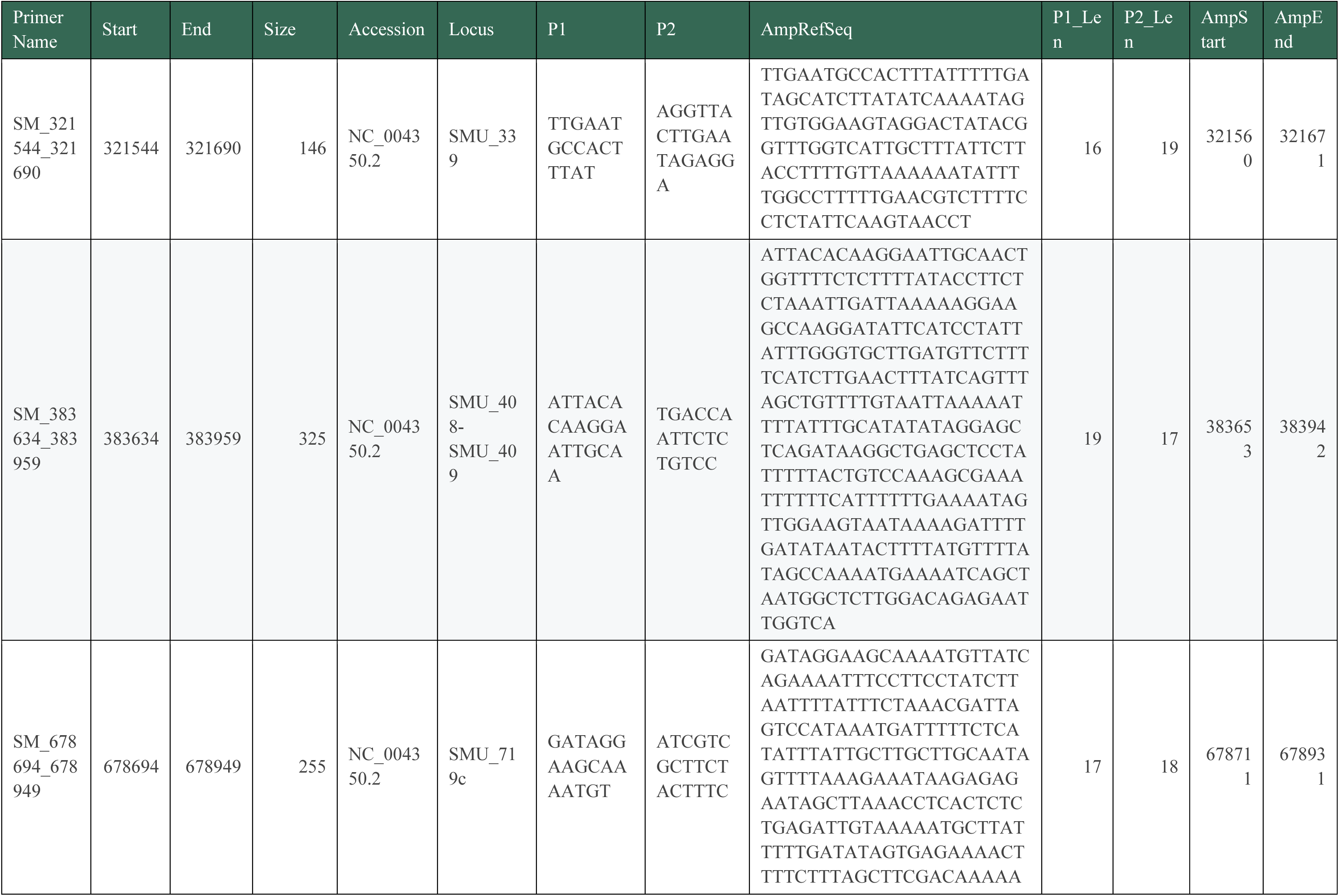
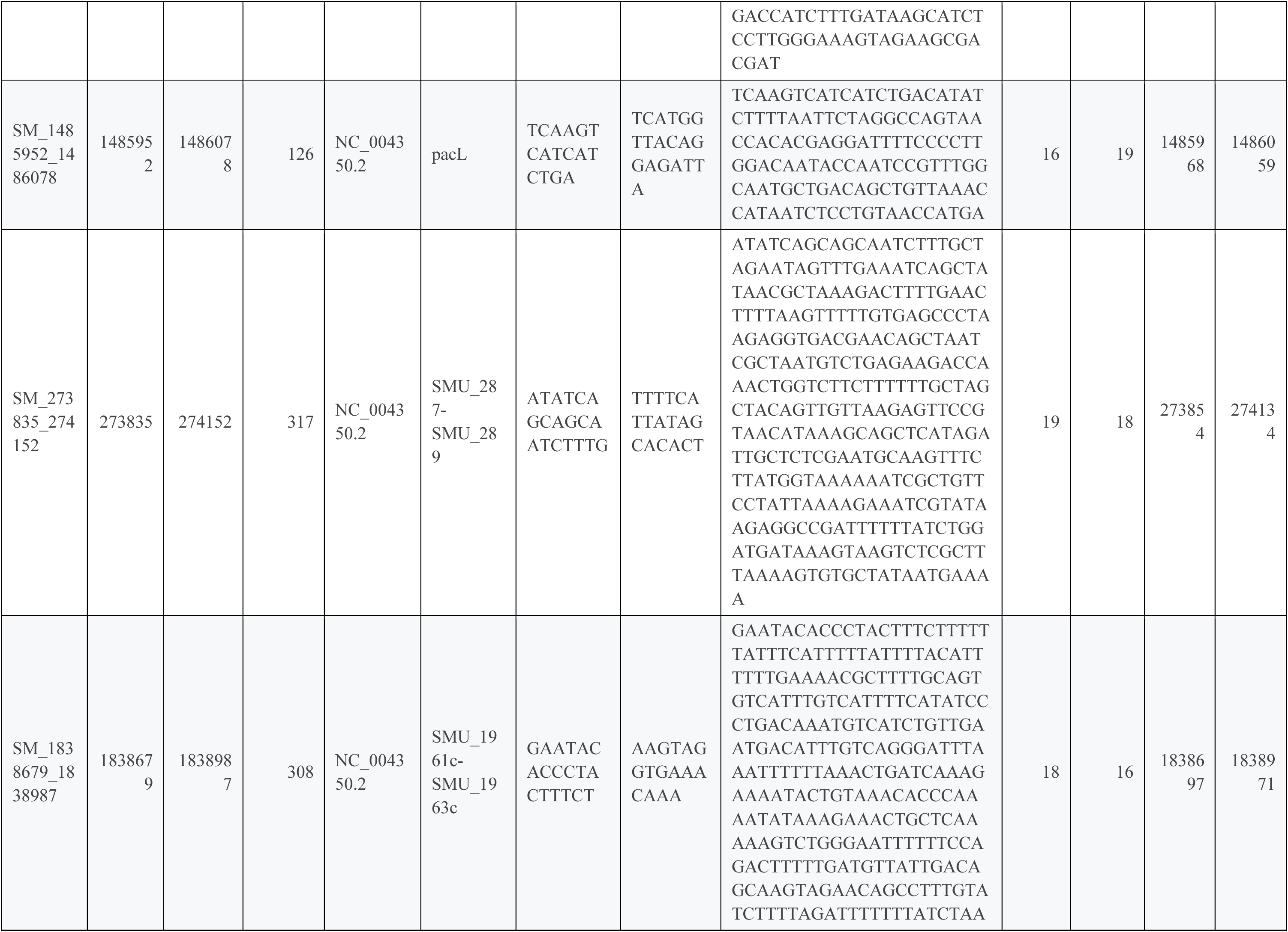
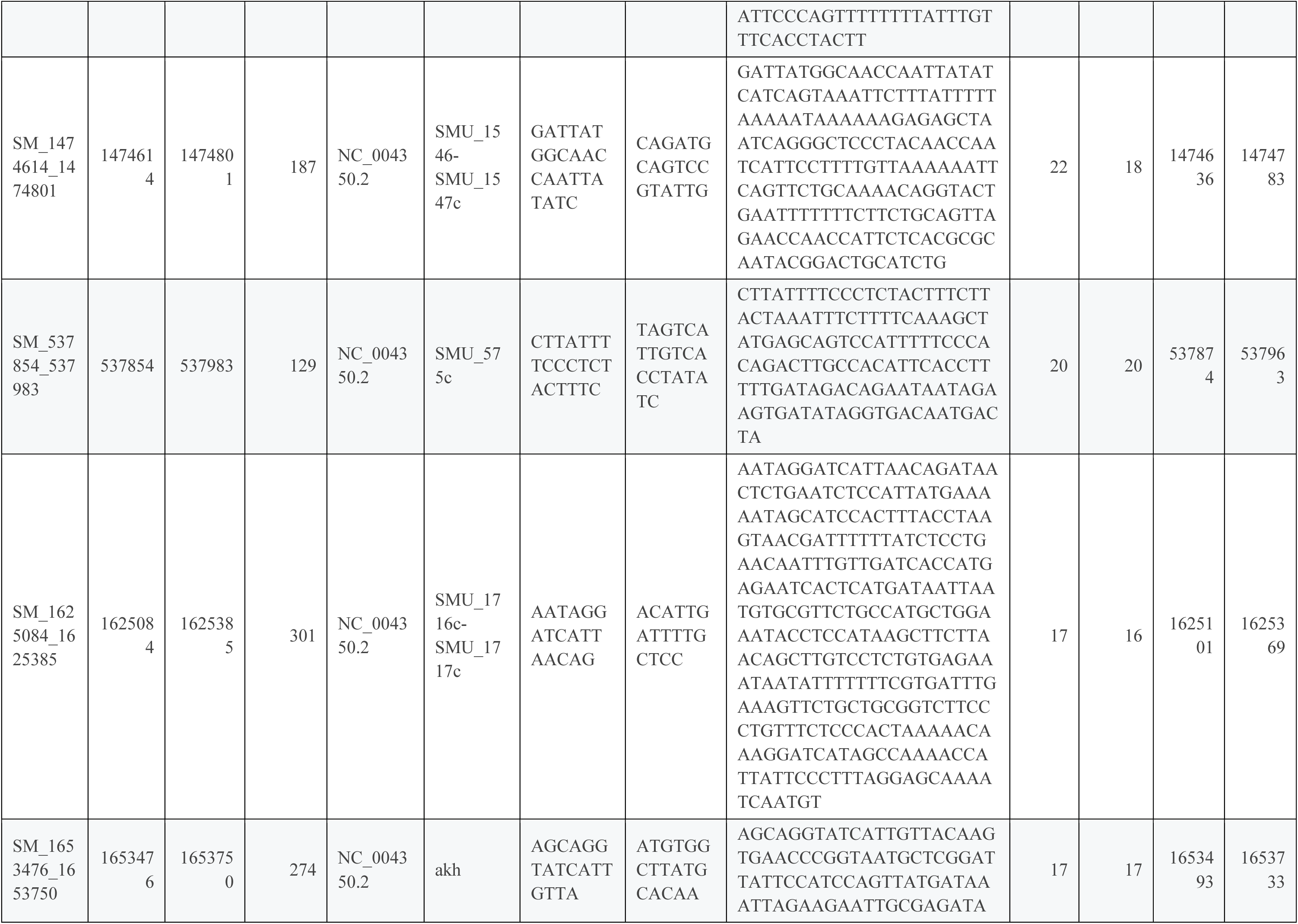
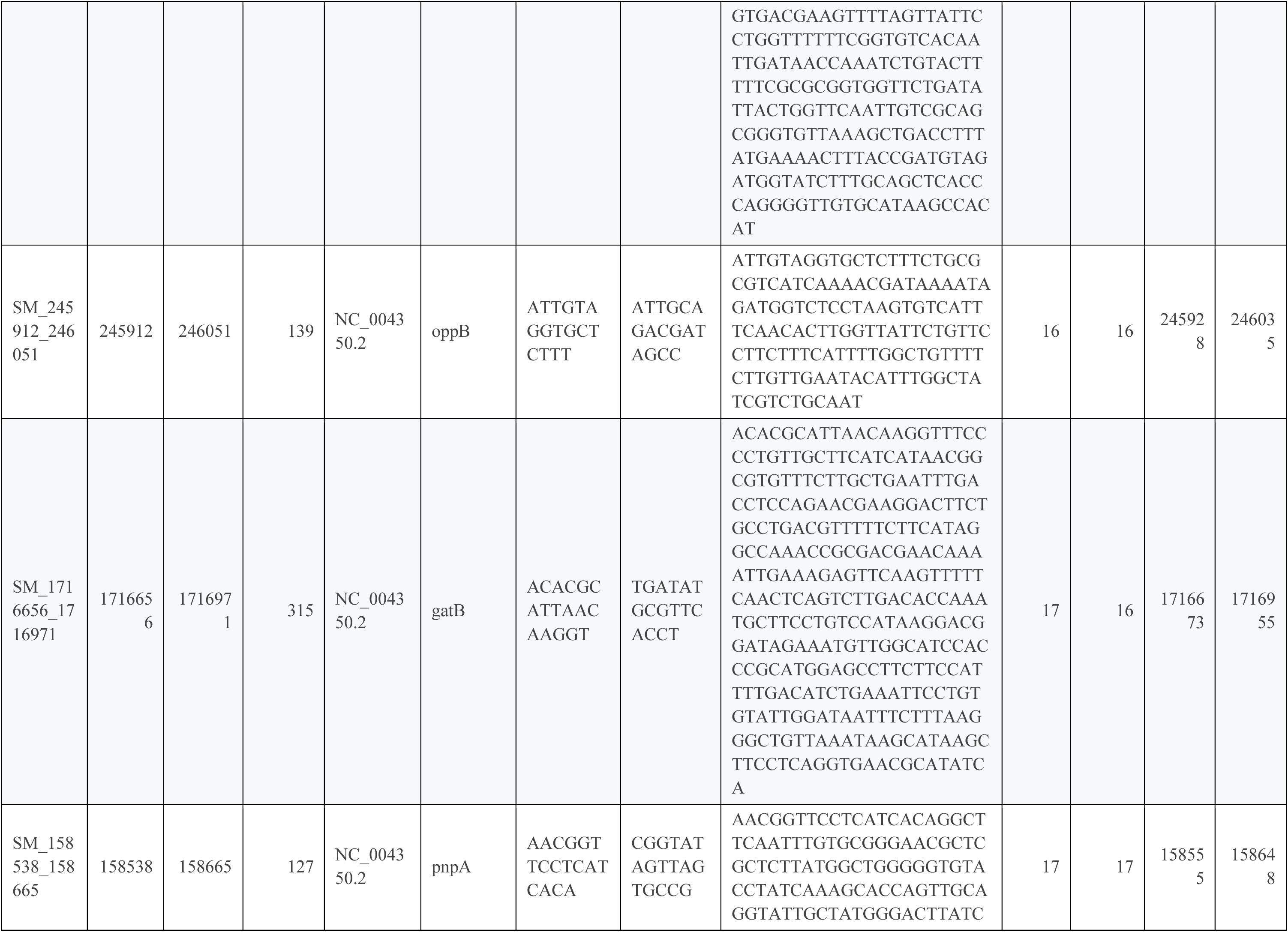
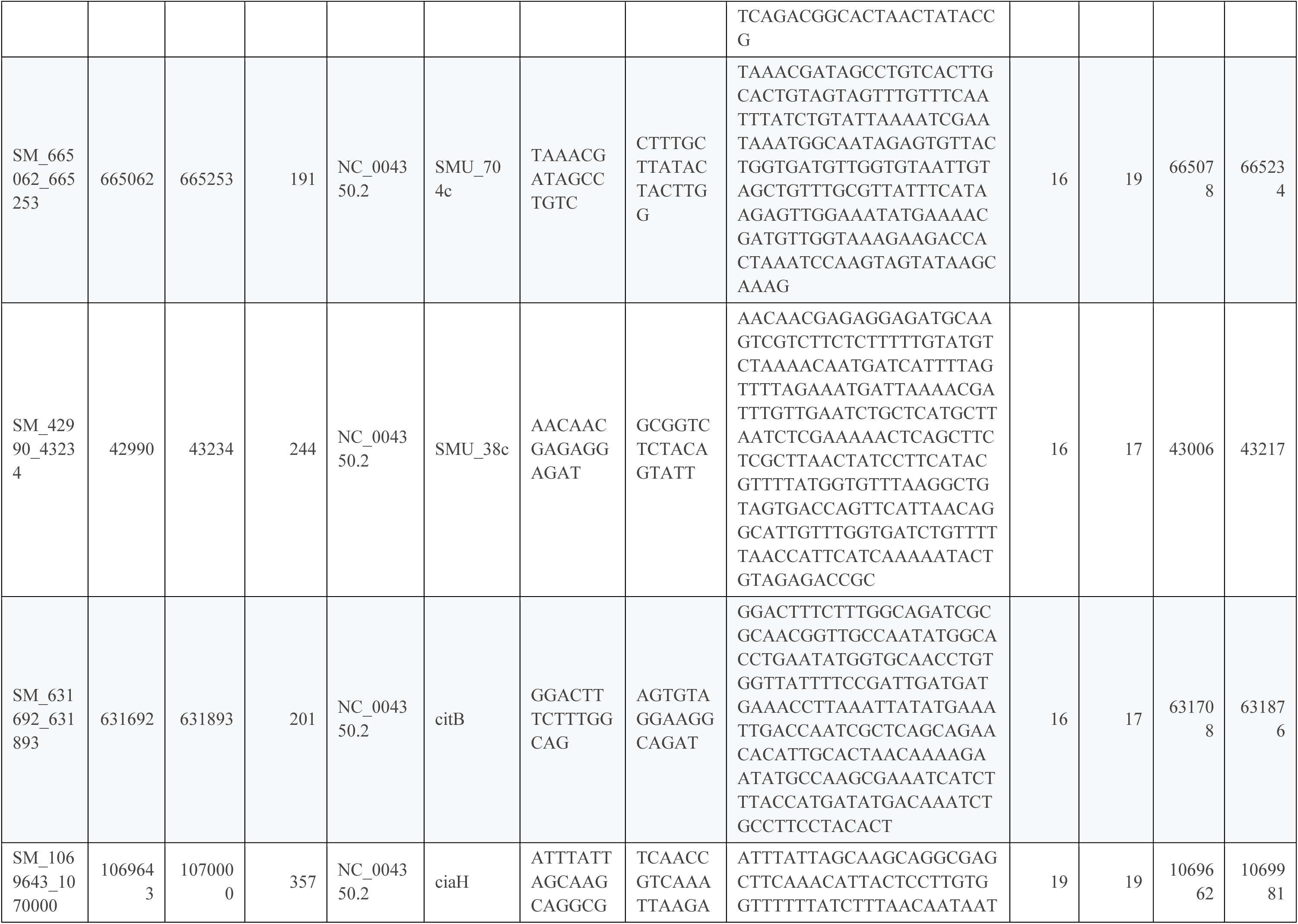
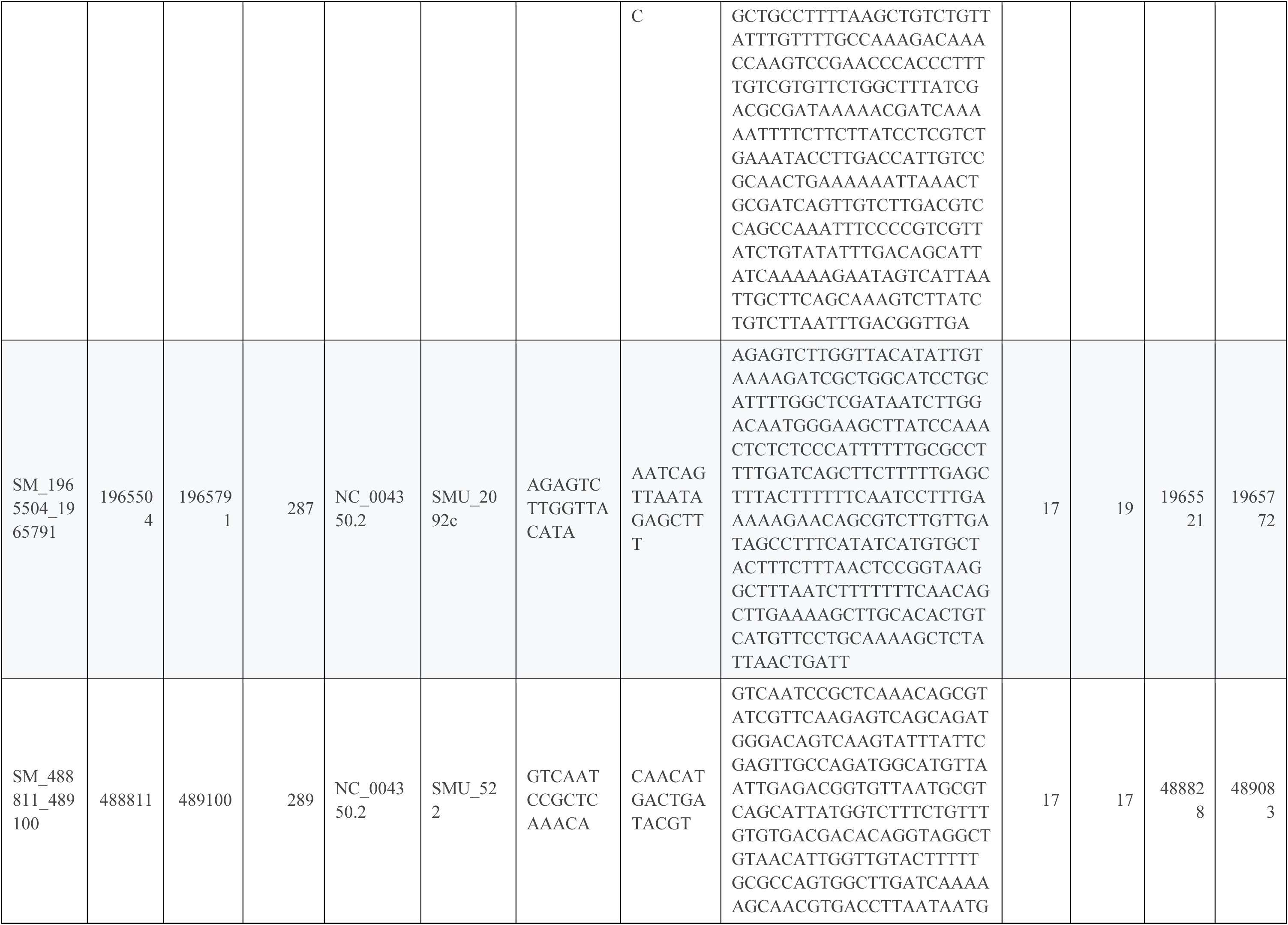
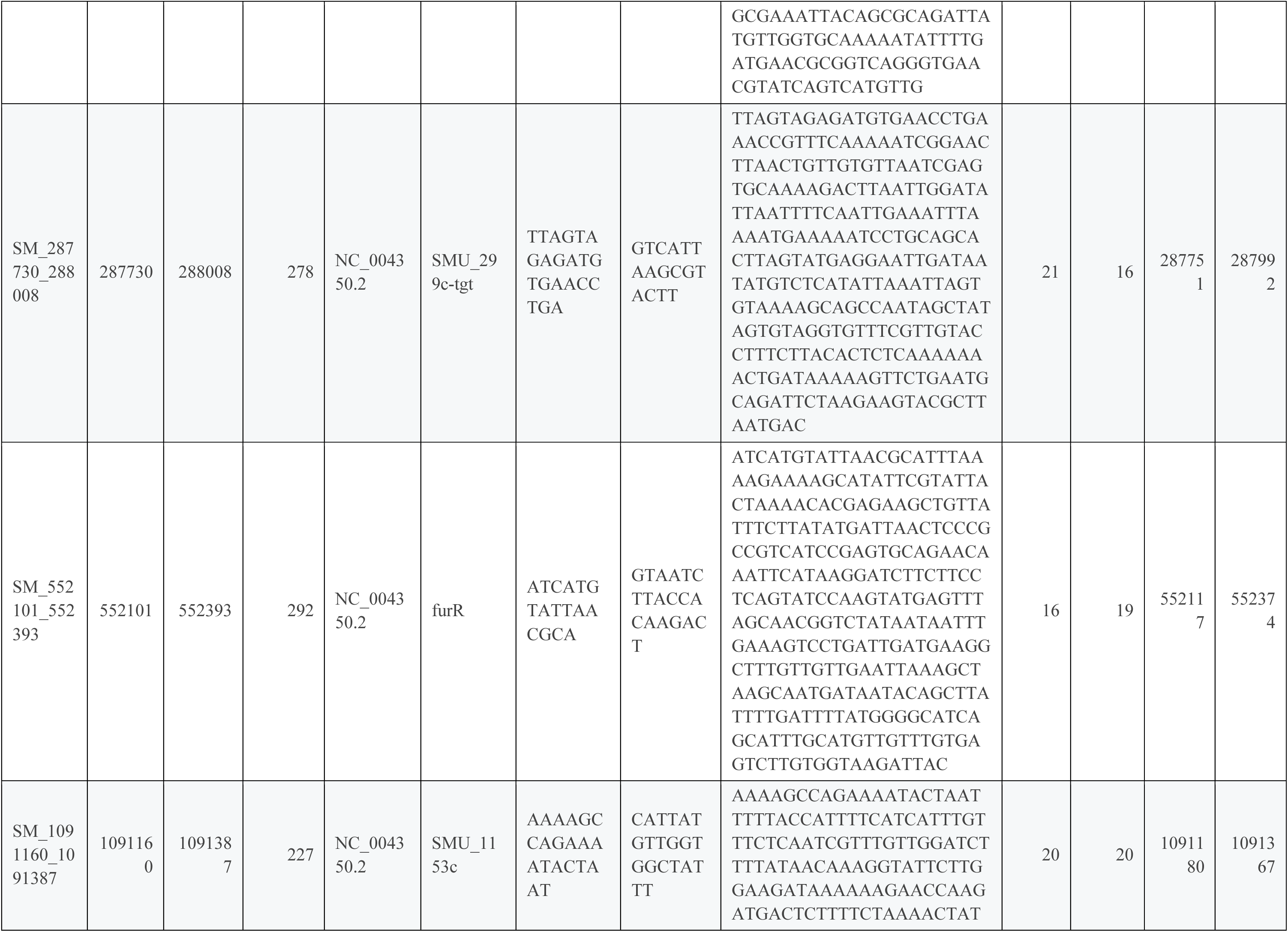
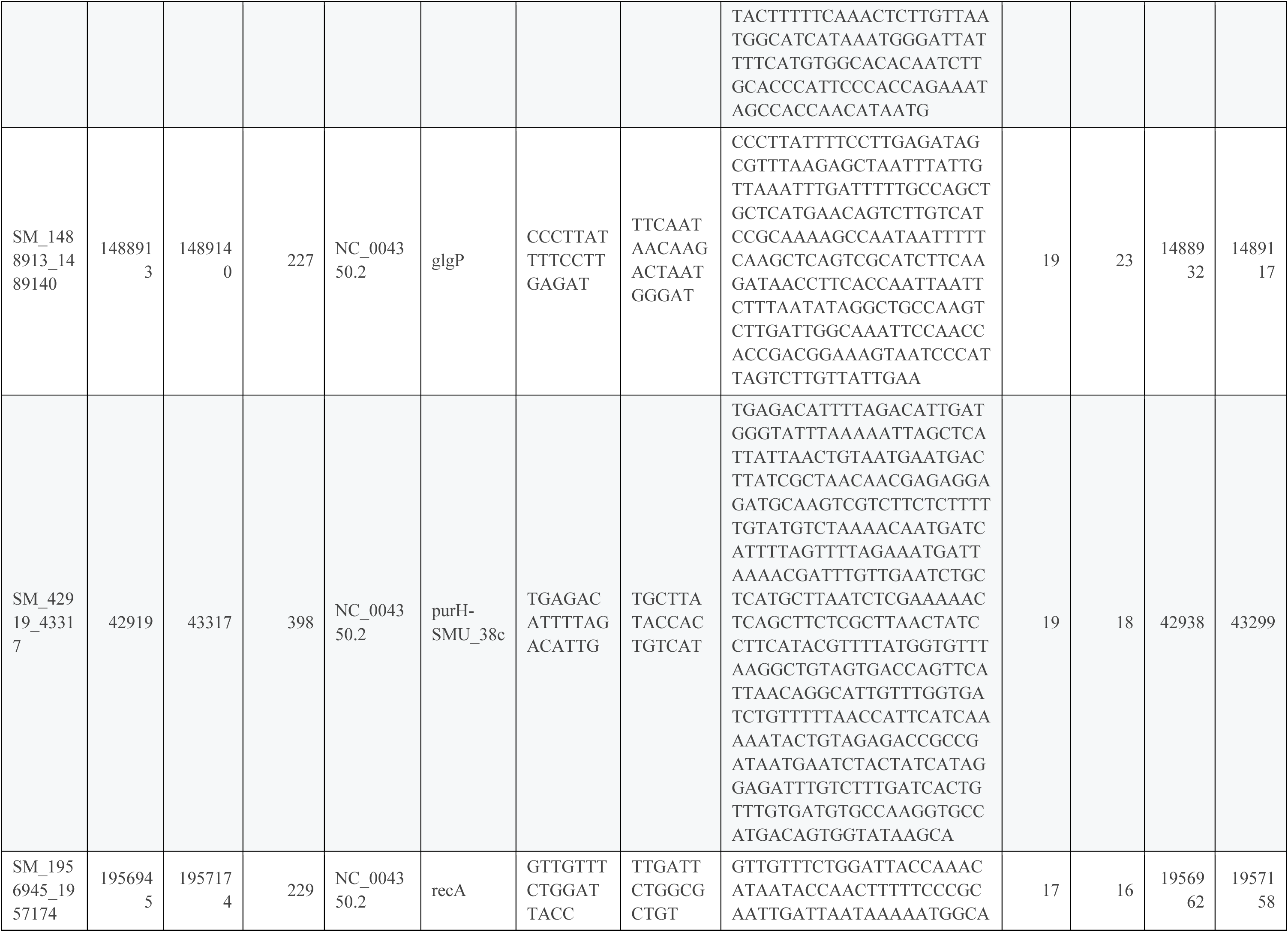
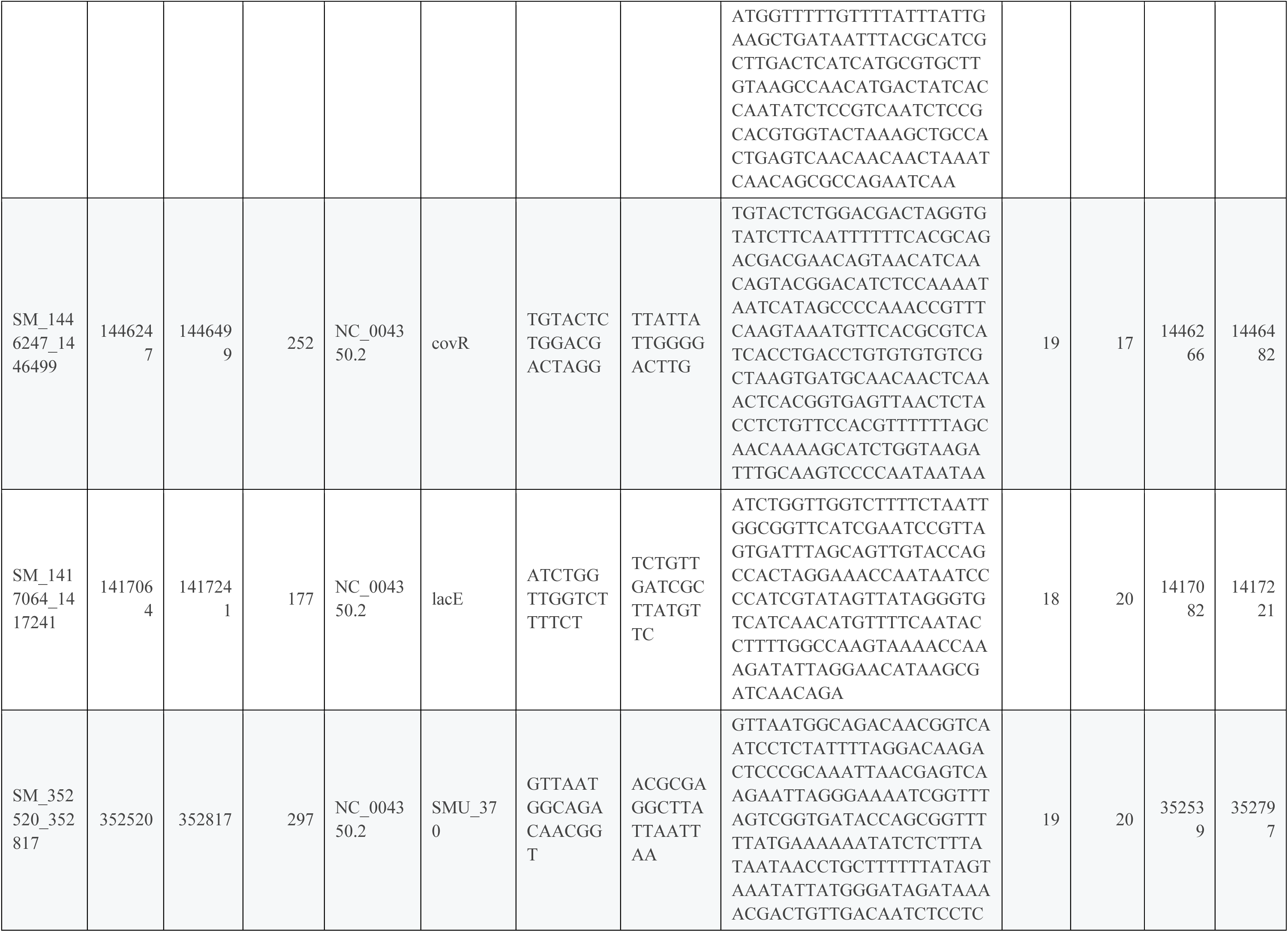
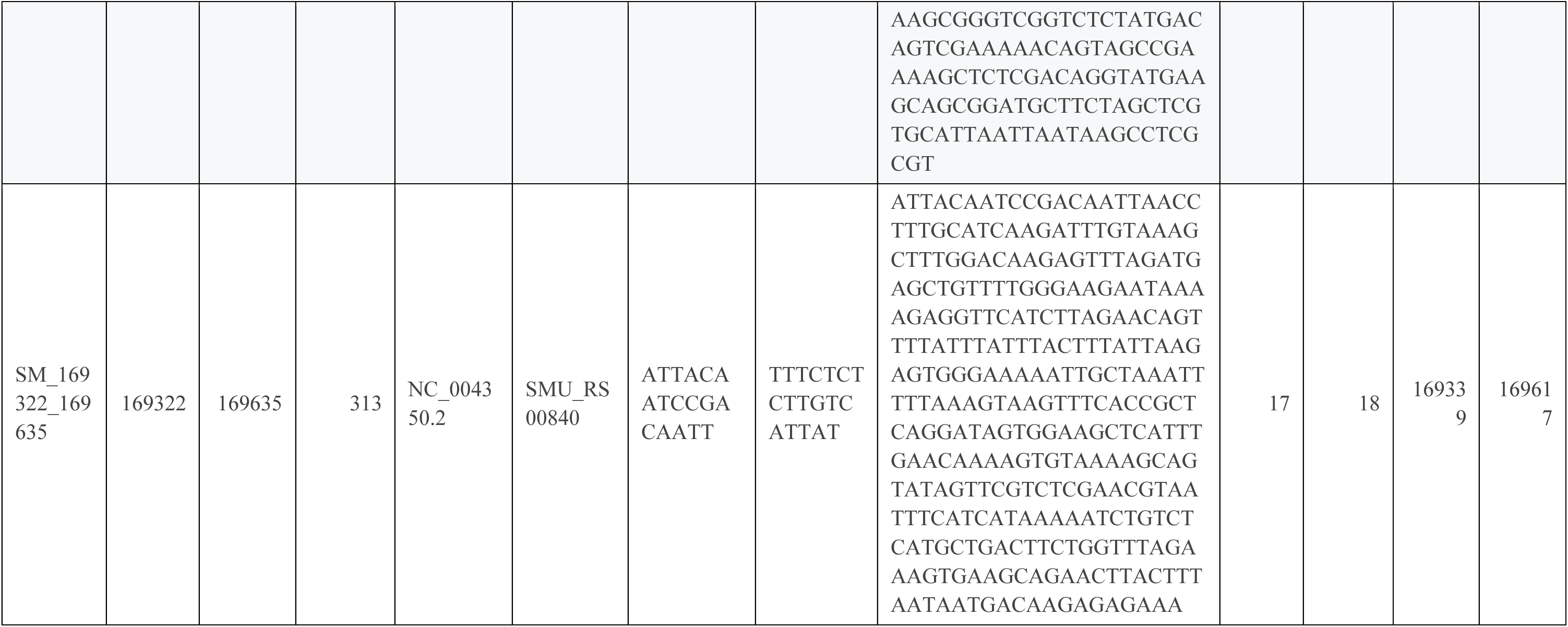
Primers used in AmpSeq assay. The primers and their positions with the reference sequences are provided along with the expected amplicon sizes in the reference genome. The locus ID is provided for amplicons within or between coding regions.

**Supplemental Table 3:**
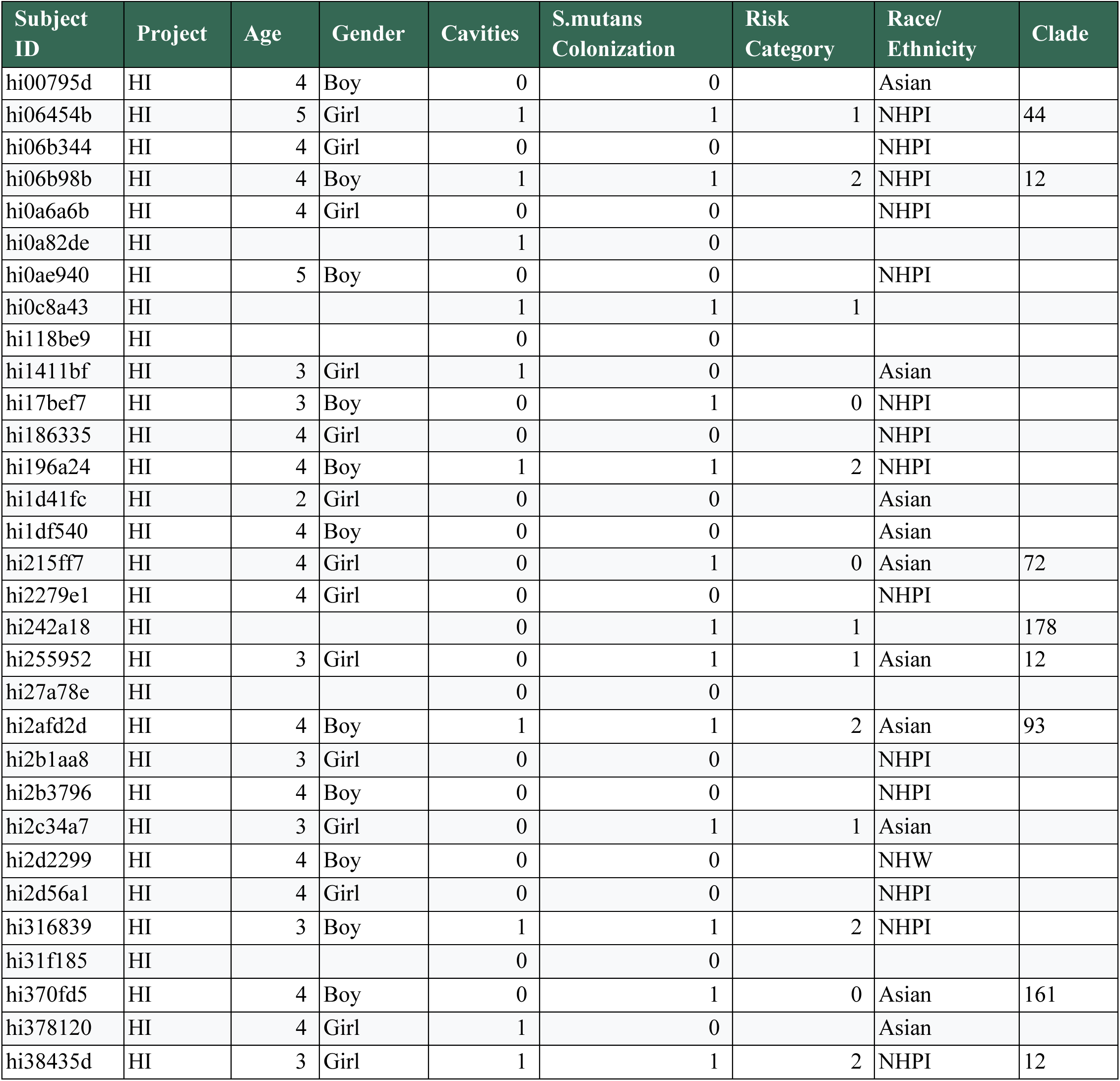
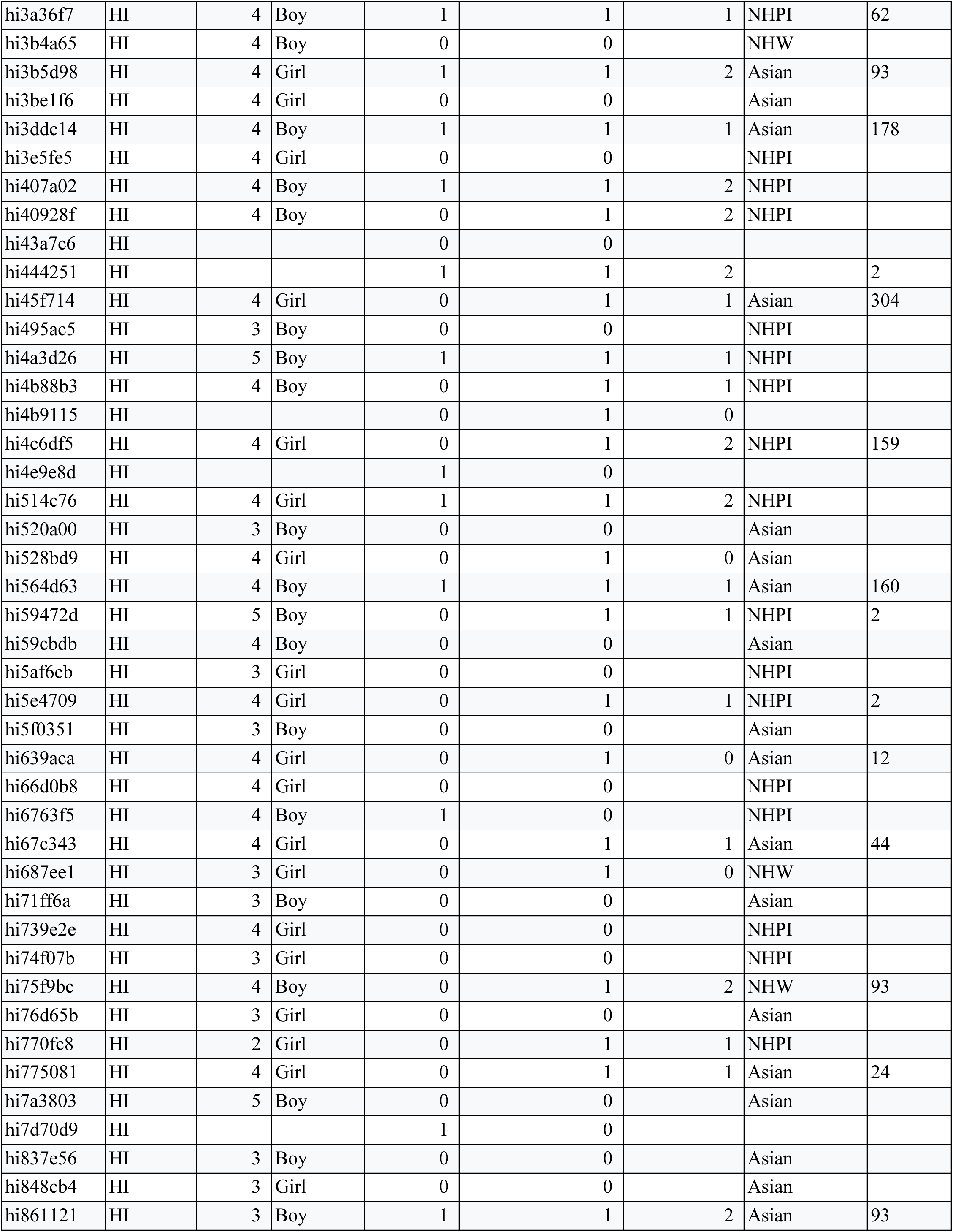
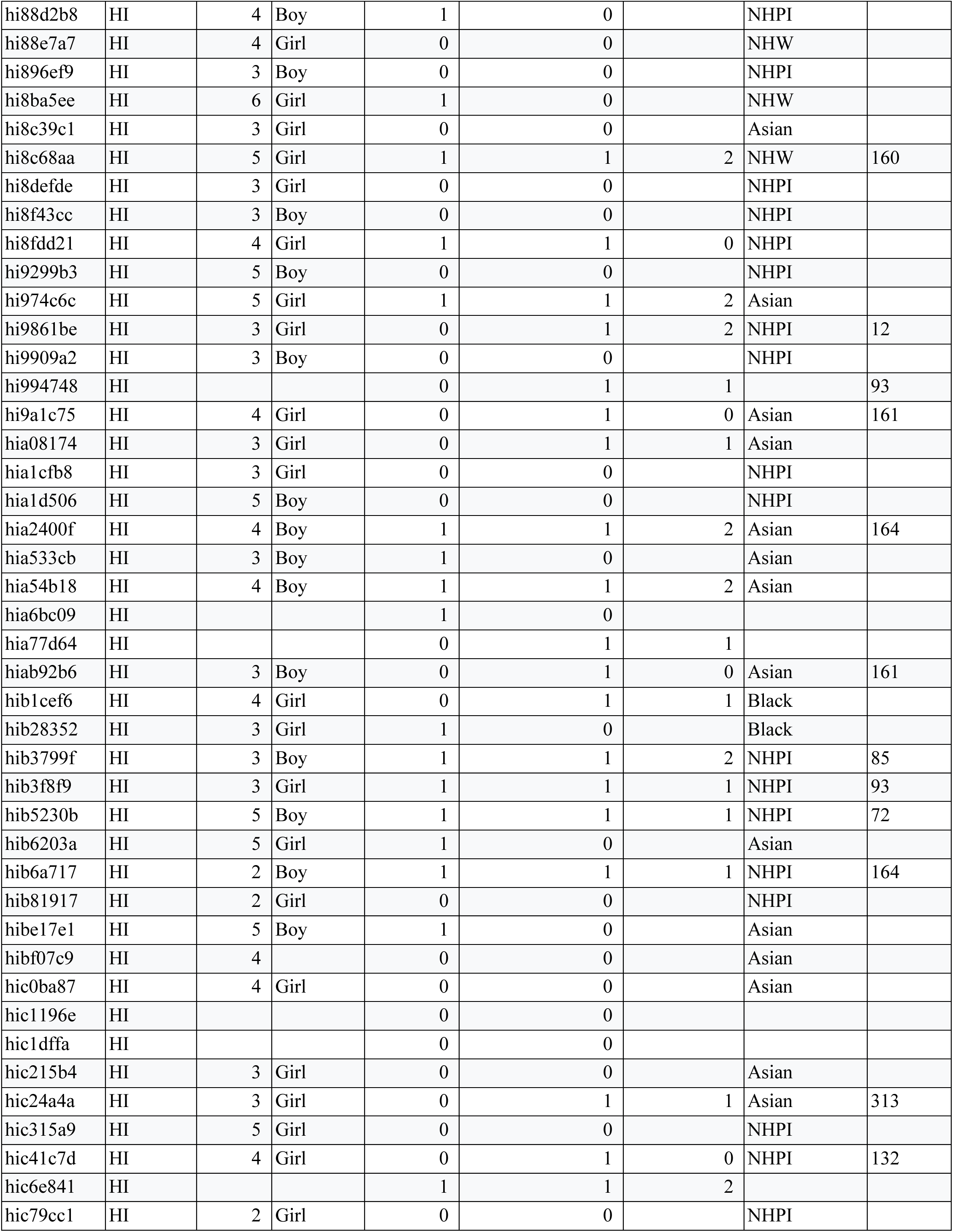
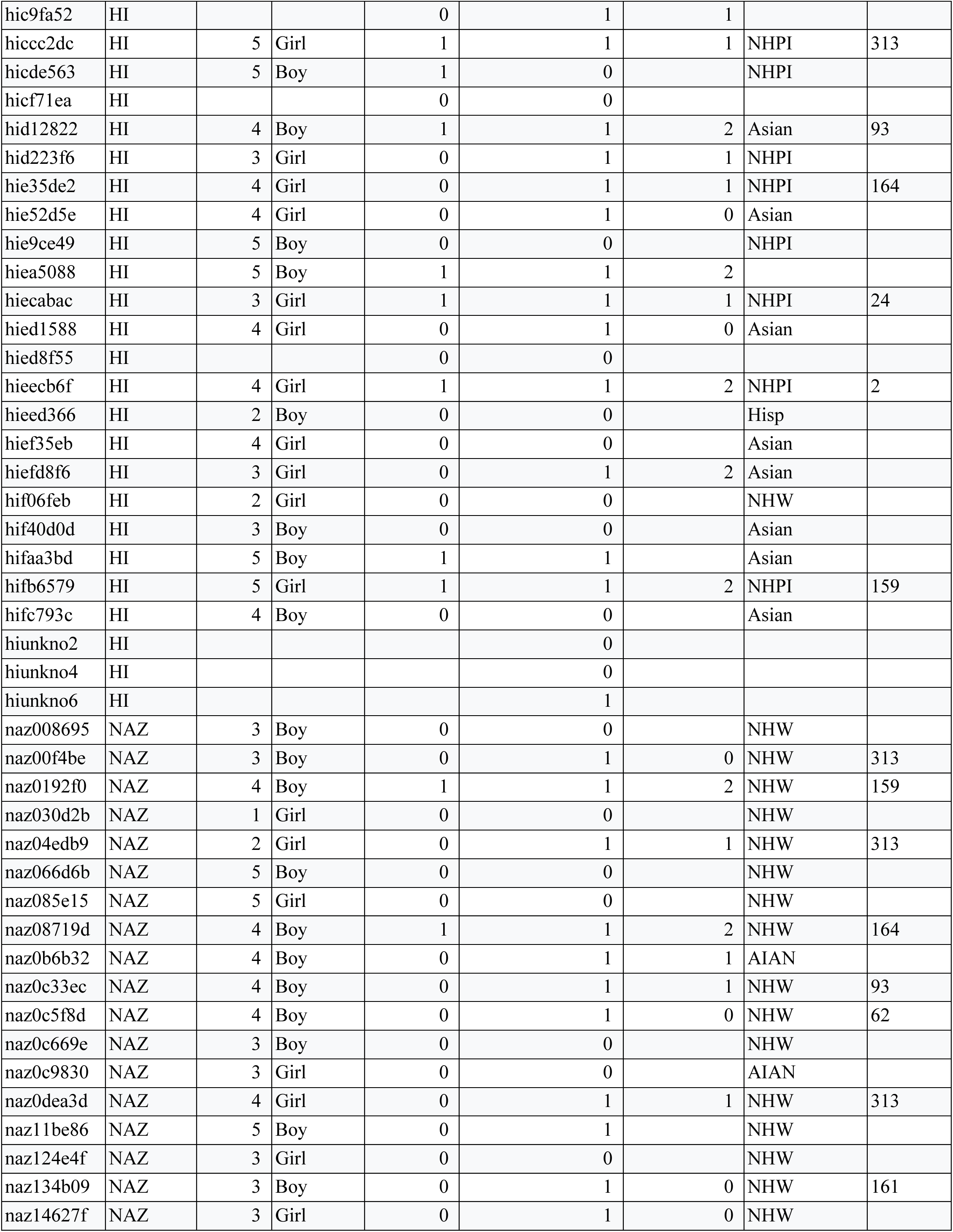
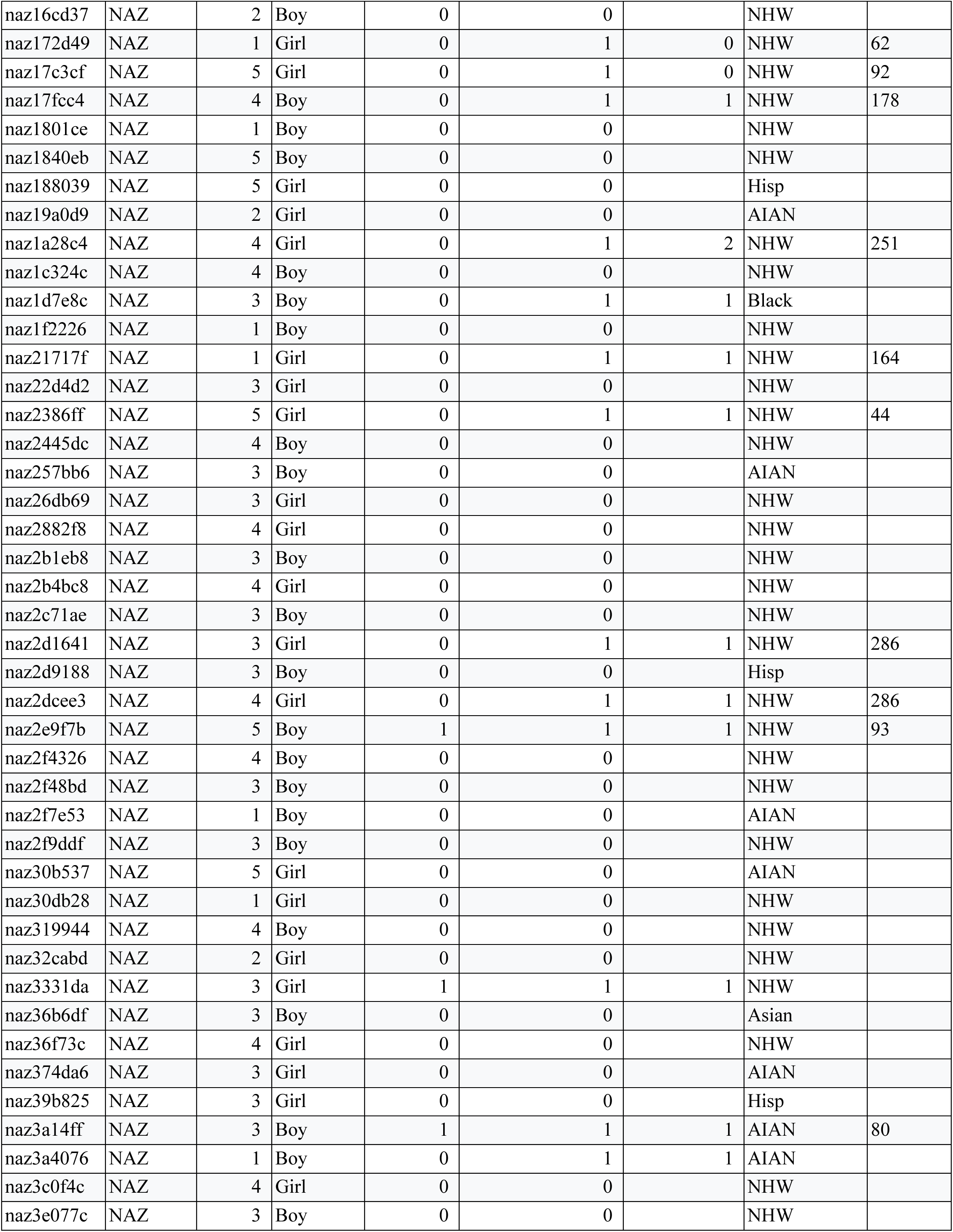
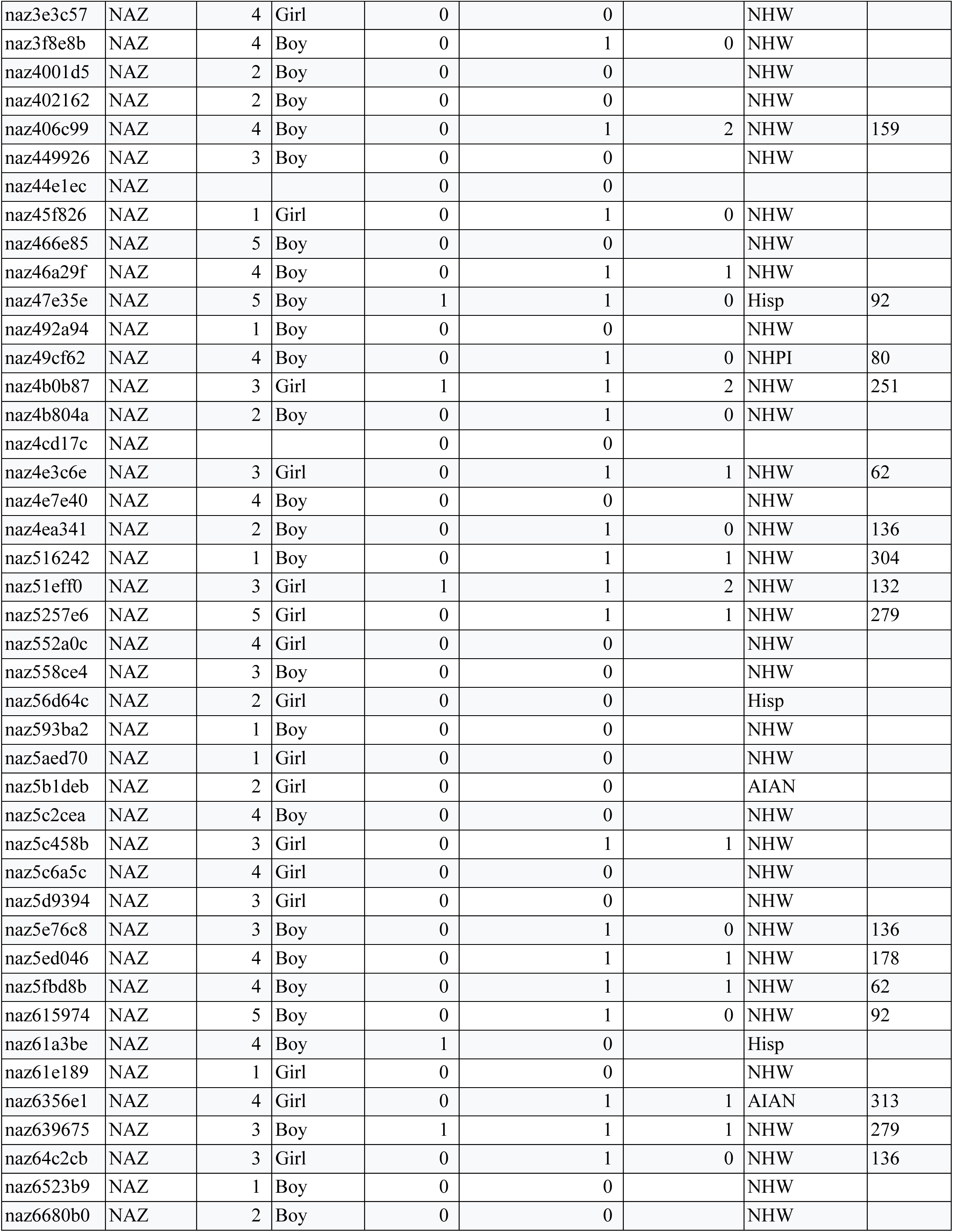
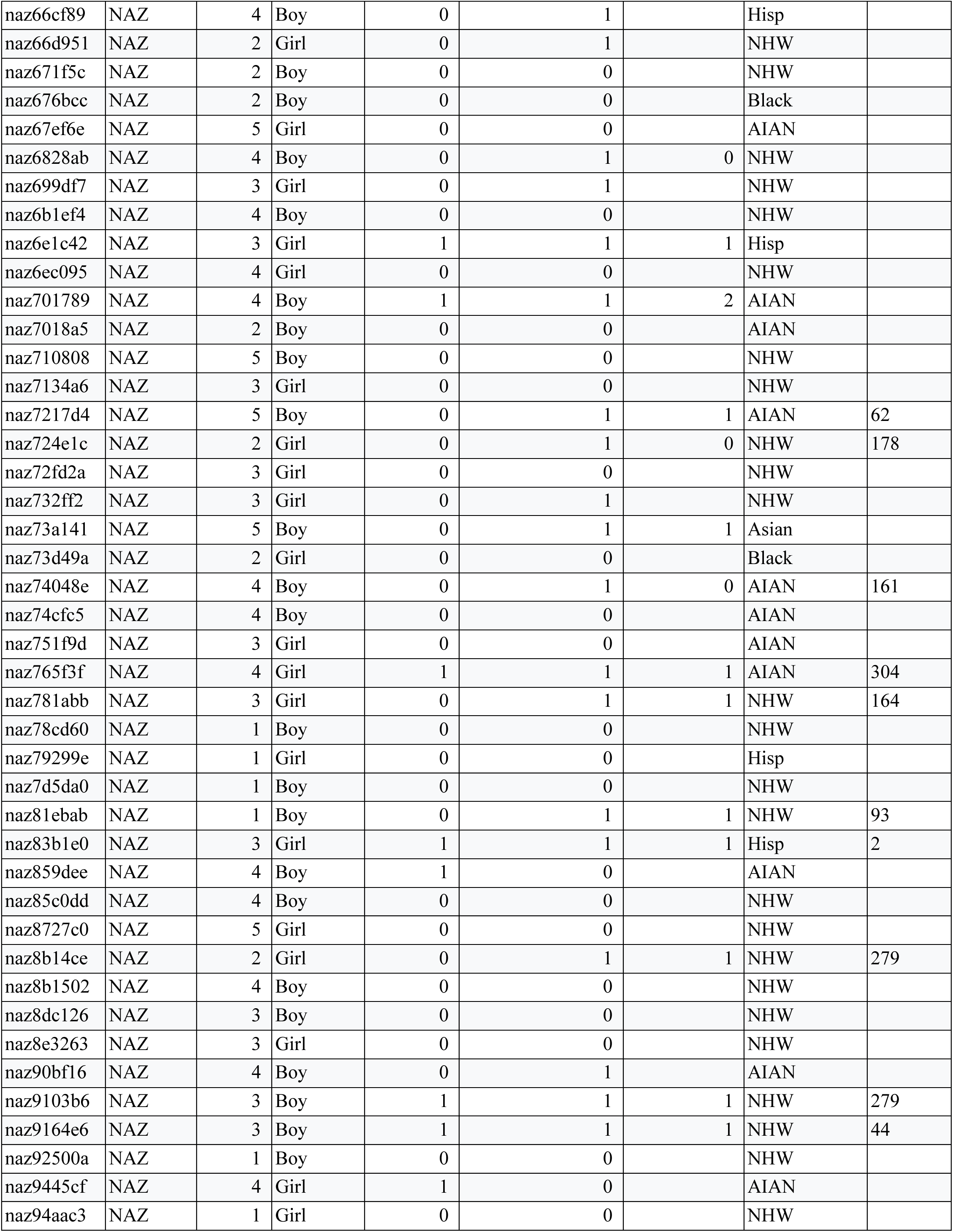
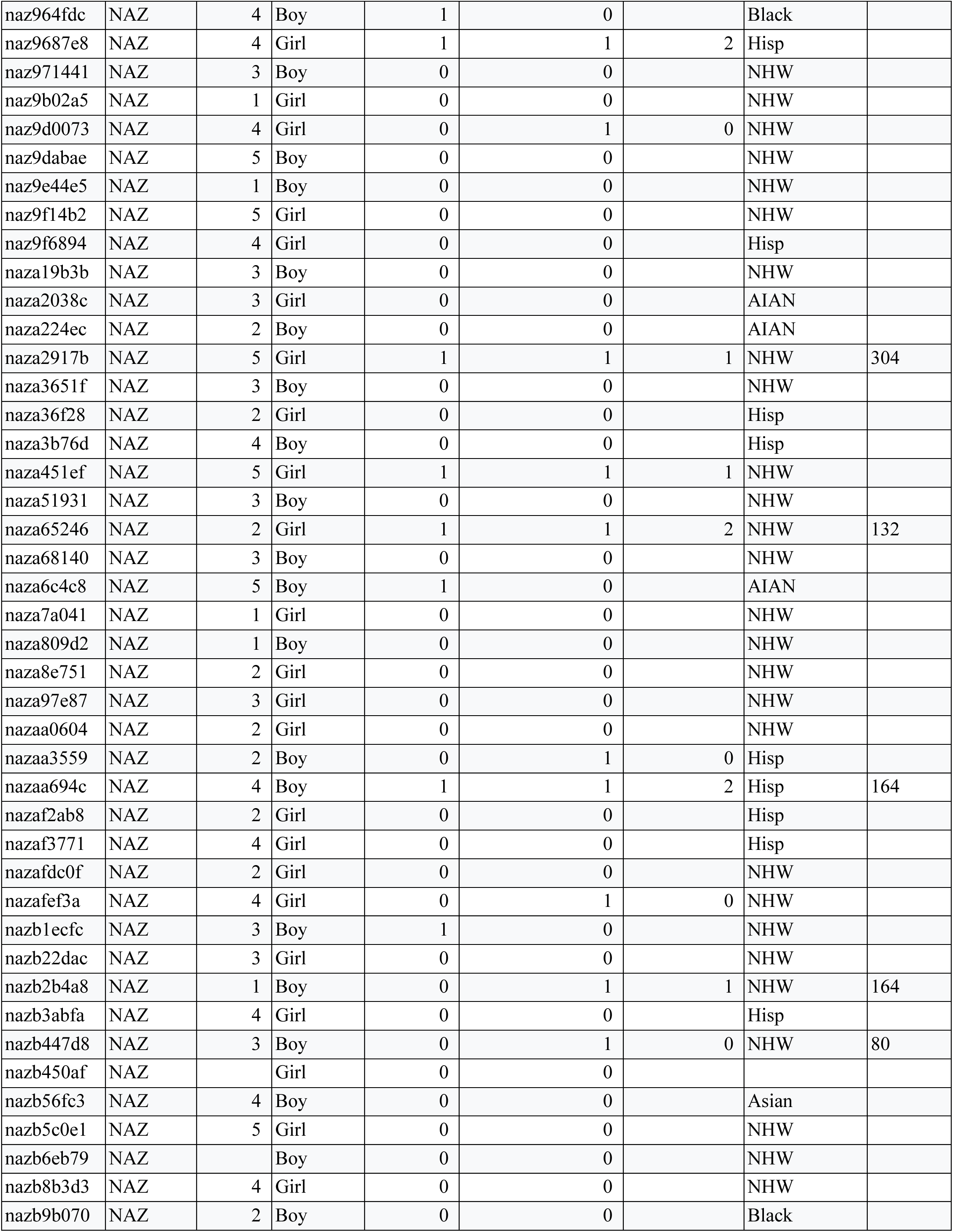
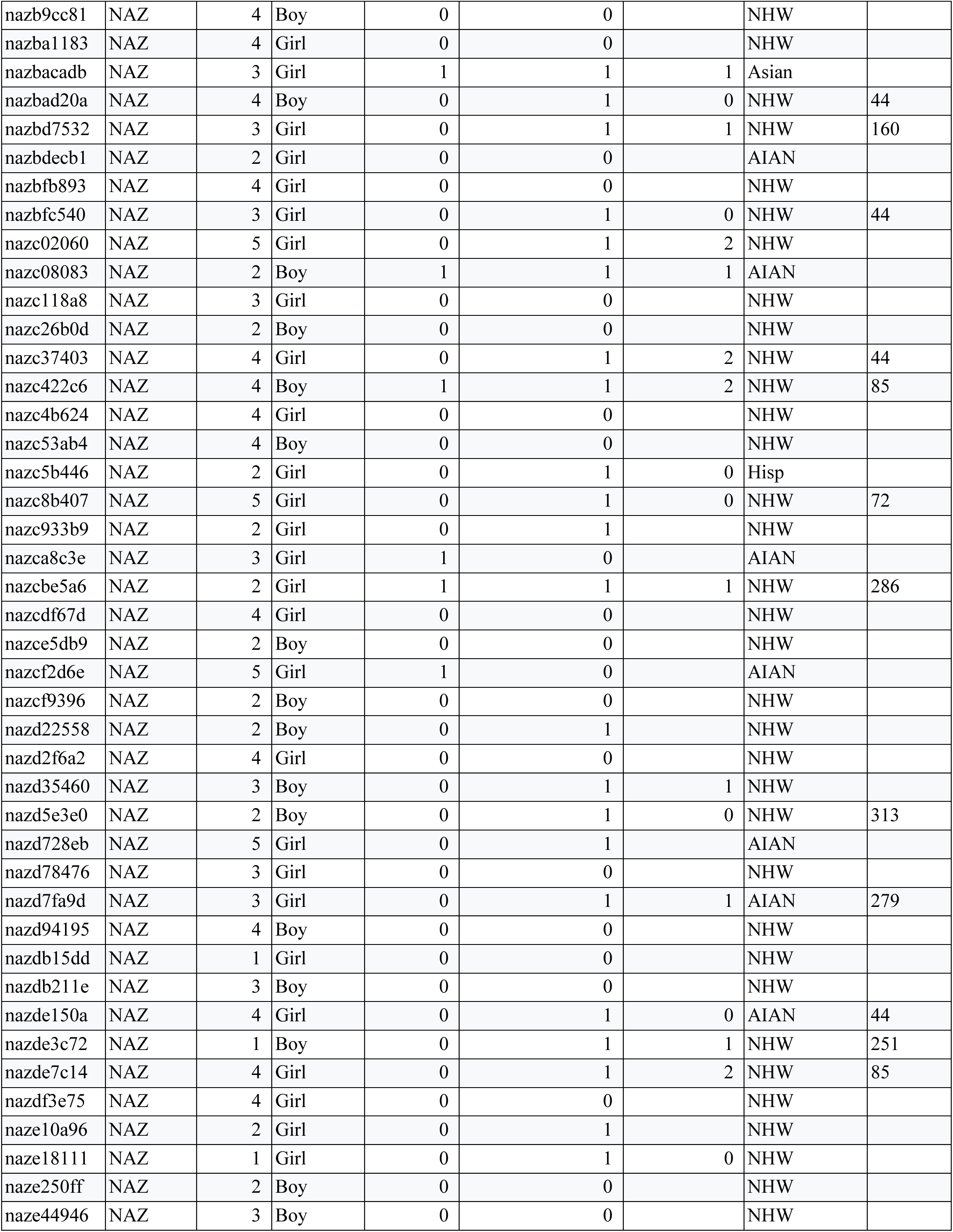
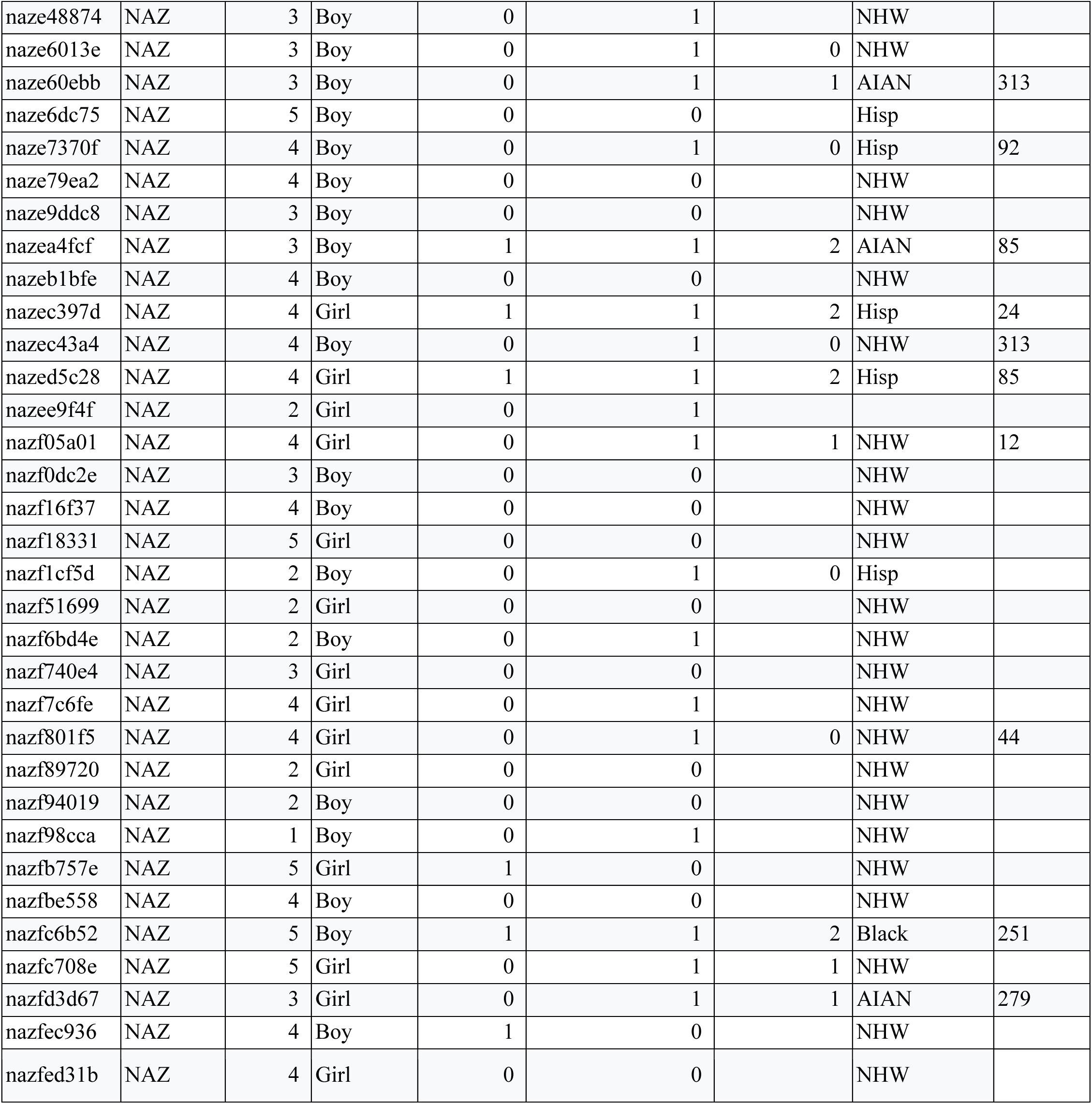
Participant metadata.

**Supplemental Table 4:** *S. mutans* core genome SNPs that PySeer identified as significant.

