## Supplementary figures and images for "Biological Drivers of Early Childhood Caries in Preschool Children of Northern Arizona and Hawaii"

### Supplemental Figure 1

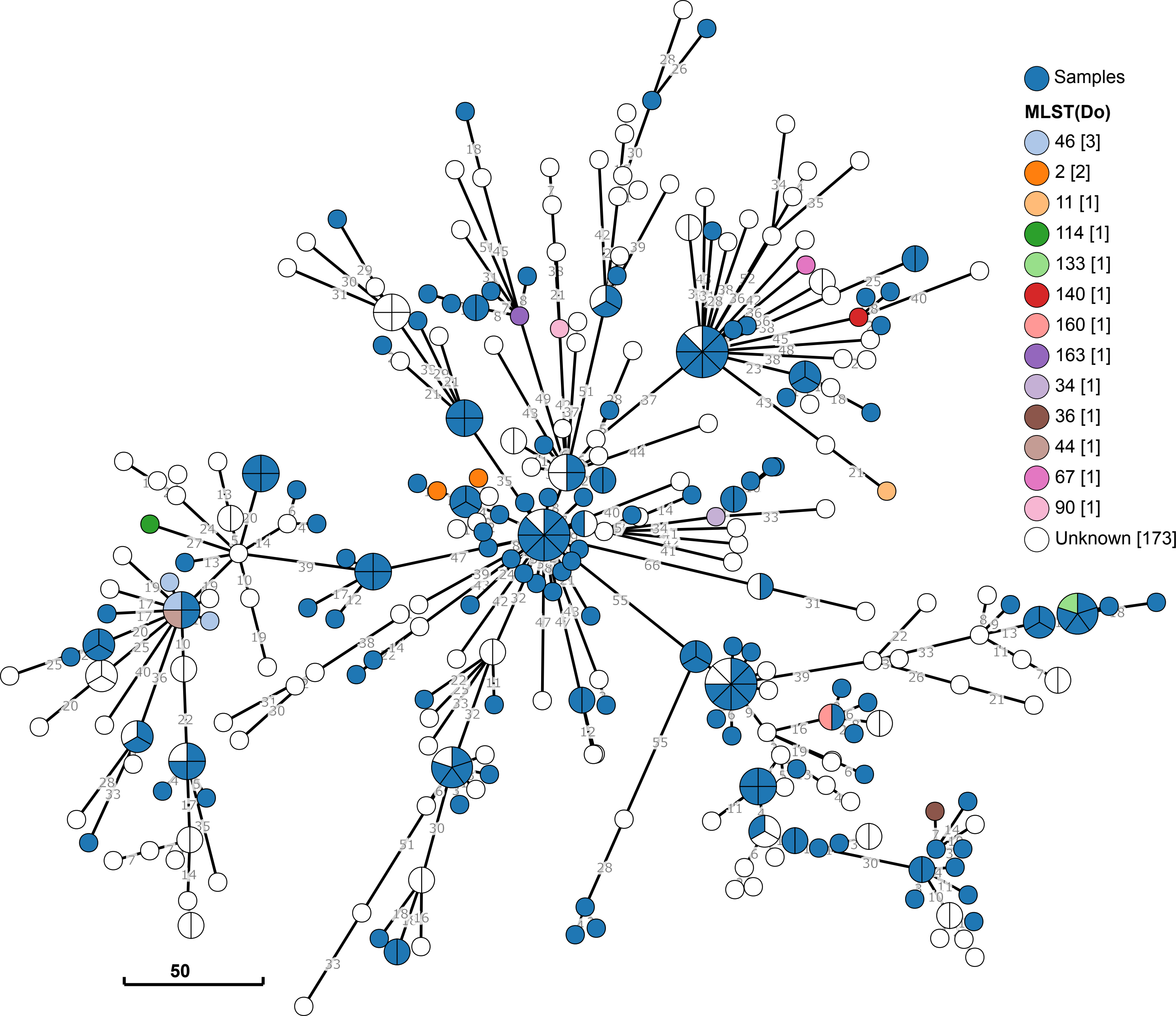

### Supplemental Figure 2

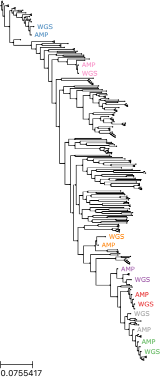

### Supplemental Figure 3

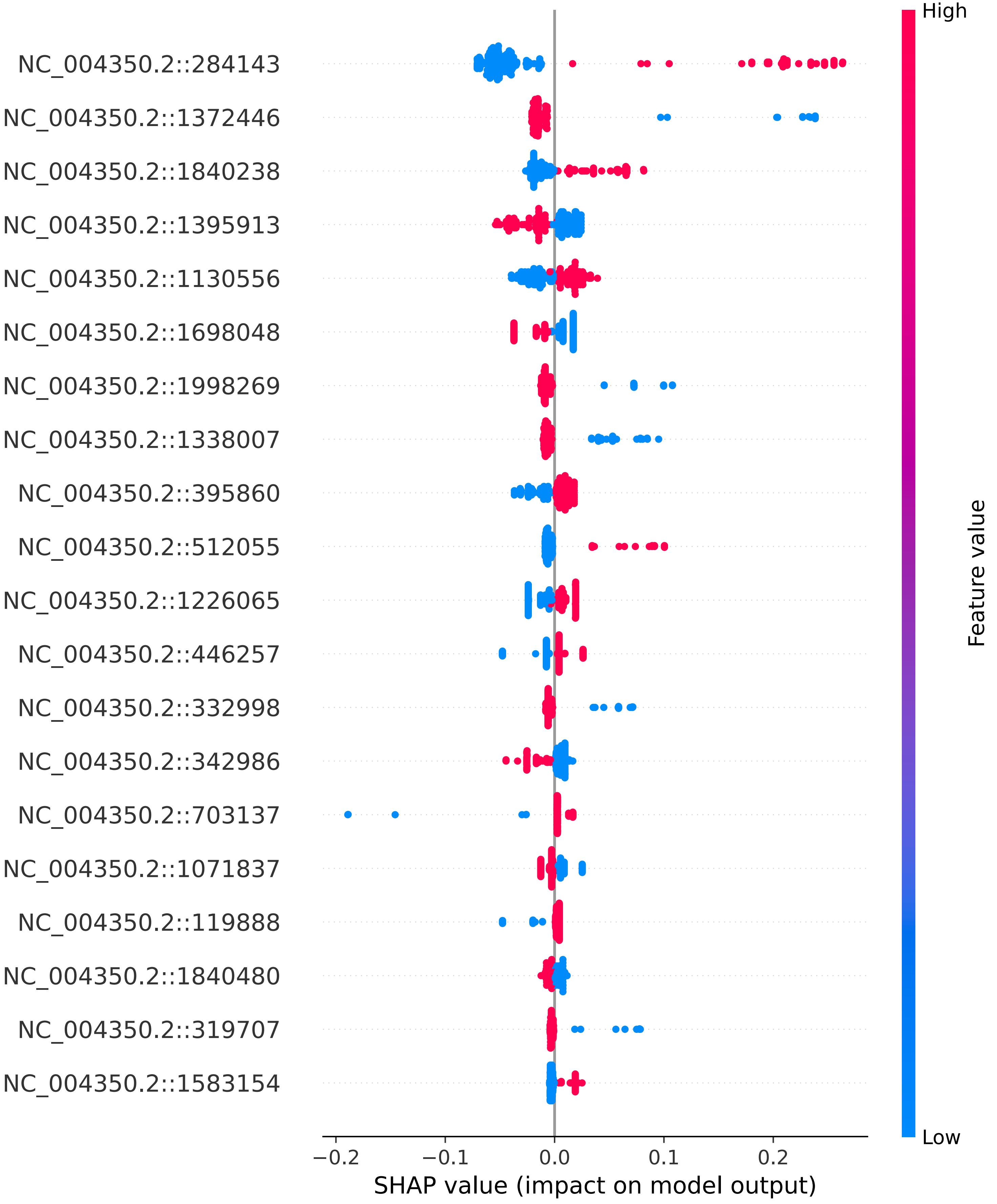

### Supplemental Figure 4

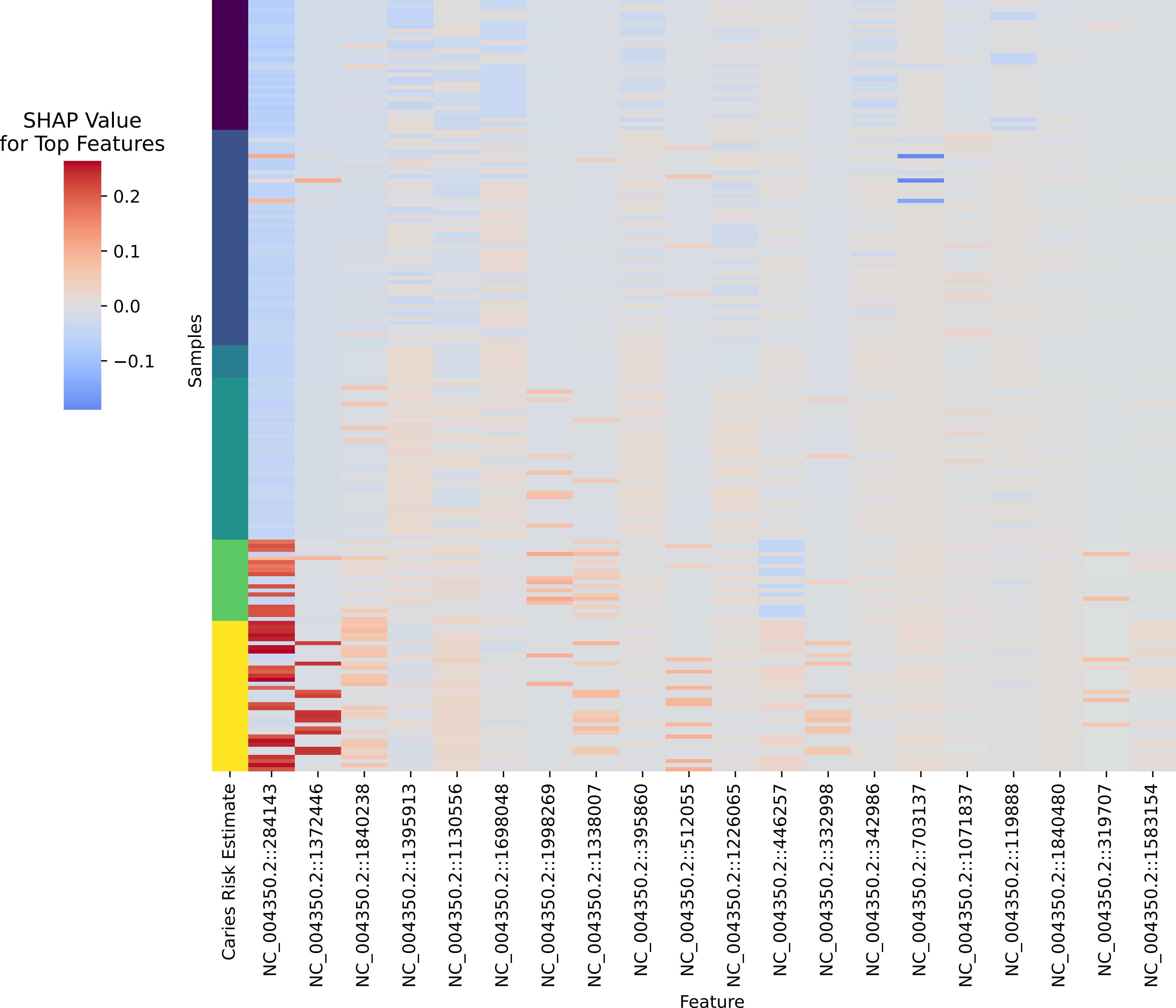
